# Coordinated membrane remodeling and fission-fusion drive mitochondrial patterning during development

**DOI:** 10.1101/2025.06.09.658059

**Authors:** H Aravind, Vivek Kumar, Vaishali Modha, Manish Jaiswal

## Abstract

How complex organelles remodel their architecture during development remains a fundamental question in cell biology. Using the *Drosophila* nebenkern, a giant multilamellar mitochondrion that transforms into two elongated mitochondrial derivatives during spermiogenesis, we dissect the challenges underlying large organelle morphogenesis. We show that this transformation is achieved through coordinated control of fusogenicity, membrane geometry, and membrane redistribution. PINK1/Parkin-dependent downregulation of the mitofusin Marf reduces mitochondrial fusogenicity and stabilizes organelle separation following division. Concurrently, membrane tubulation establishes a geometric template for DRP1 recruitment, specifying a deterministic fission plane within a large mitochondrial sheet structure. Mitochondrial elongation is accompanied by the formation of mitochondria-derived inverted vesicles (MDiVs), previously undescribed double-membraned structures that may redistribute membrane through inward budding independently of canonical fission and fusion. More broadly, our work establishes a framework for understanding how membrane geometry, regulated fusogenicity, and vesicle-mediated remodeling cooperate to generate complex organelle architectures during development.

**Significance statement:** Large organelles pose unique challenges for scaling and division because canonical membrane dynamics must be coordinated across extensive membrane architectures. Using a giant multilamellar mitochondrial structure as a model, we show that organelle morphogenesis is achieved through independent yet coordinately regulated mechanisms that spatially pattern division, stabilize partitioning, and redistribute membranes through intraorganellar vesicles with an inverted topology. These findings reveal principles of organelle morphogenesis that extend beyond conventional fission-fusion models.

## Introduction

Mitochondria display remarkable morphological versatility, ranging from tubules, spheres, donuts, and lassos, both as isolated and interconnected networks, which vary across cell types, tissues, and developmental stages (1). Mitochondrial remodeling influences cellular metabolism, cell signaling, and organelle quality control, thereby shaping cellular processes such as cell division (2, 3), differentiation (4–7), and morphogenesis. Indeed, disruptions in mitochondrial morphology have been linked to various diseases and affect several developmental processes (8, 9), underscoring the need to understand mechanisms that regulate mitochondrial shape.

Mitochondrial architecture is governed by a coordinated cycle of fission and fusion, mediated primarily by conserved small GTPases Mitofusins (Marf in *Drosophila*), OPA1, and Drp1 (10). While fission and fusion dynamics can remodel mitochondrial size and connectivity, they cannot fully account for the diverse ultrastructural forms observed *in vivo* (*11*). As recently found, inner membrane rearrangement via cristae remodeling can drive mitochondrial elongation (12) and regulate unique morphological changes such as mitochondrial pearling (13), suggesting that additional layers of membrane remodeling, together with canonical fission-fusion, drive mitochondrial shape transitions. How these diverse determinants of mitochondrial shape interplay to regulate mitochondrial morphology during complex developmental transitions remains underexplored.

Scaling of mitochondrial size remains an important property through which cells regulate mitochondrial positioning, activity, and repair. For example, during stress, mitochondria undergo hyperfusion, changing their network characteristics to evade apoptotic stress and mitophagy (14, 15). During oogenesis, they undergo extensive fission to aid quality control (16). Accordingly, morphogenetic events, such as the establishment of epithelial apical polarity (17), neuronal development (18), and spermiogenesis (19), require highly coordinated changes in mitochondrial shape, size, localization, and function (9). During insect spermiogenesis, post-meiotic mitochondrial size scales over several orders of magnitude to form what is perhaps the most complex mitochondrial structure, namely the nebenkern. Nebenkern mitochondria display a spherical, whorl-like arrangement that later bifurcates into two distinct mitochondrial derivatives, a process crucial for sperm elongation and morphogenesis and thereby fertility (20). Systematic mitochondrial shape transitions during spermiogenesis thus make it a valuable model system to investigate how mitochondrial scaling is coordinated during development.

Here, using high-resolution and expansion microscopy to capture mitochondrial shape transition during *Drosophila* spermiogenesis, we uncover a developmentally programmed switch in mitochondrial dynamics that couples reduced fusion competence with spatially confined, geometry-licensed fission, and membrane remodeling to drive nebenkern bifurcation and elongation. Moreover, we identify previously unrecognized intramitochondrial vesicular structures that may facilitate membrane redistribution during mitochondrial elongation. Together, our findings reveal how coordinated fission, fusion, and membrane remodeling drive mitochondrial shape transitions, providing a conceptual framework for understanding mitochondrial patterning across diverse contexts.

## Results

### Spermiogenesis reveals stereotypic, stage-specific mitochondrial remodeling

During spermiogenesis, mitochondria undergo sequential, stereotyped transitions in a stage-specific manner. Confocal imaging of whole-mount testis, TOM20::mCherry to label mitochondria, captures distinct morphological states across developmental stages **(Fig. 1A).** While these broad features have been described (20, 21), their underlying ultrastructural organization remains incompletely resolved. To address this, we combined expansion microscopy with SoRa super-resolution imaging to achieve high-resolution volumetric visualization of mitochondrial architecture across stages. This approach revealed previously unresolved structural details, enabling a precise characterization of mitochondrial remodeling during spermiogenesis.

**Fig 1:**
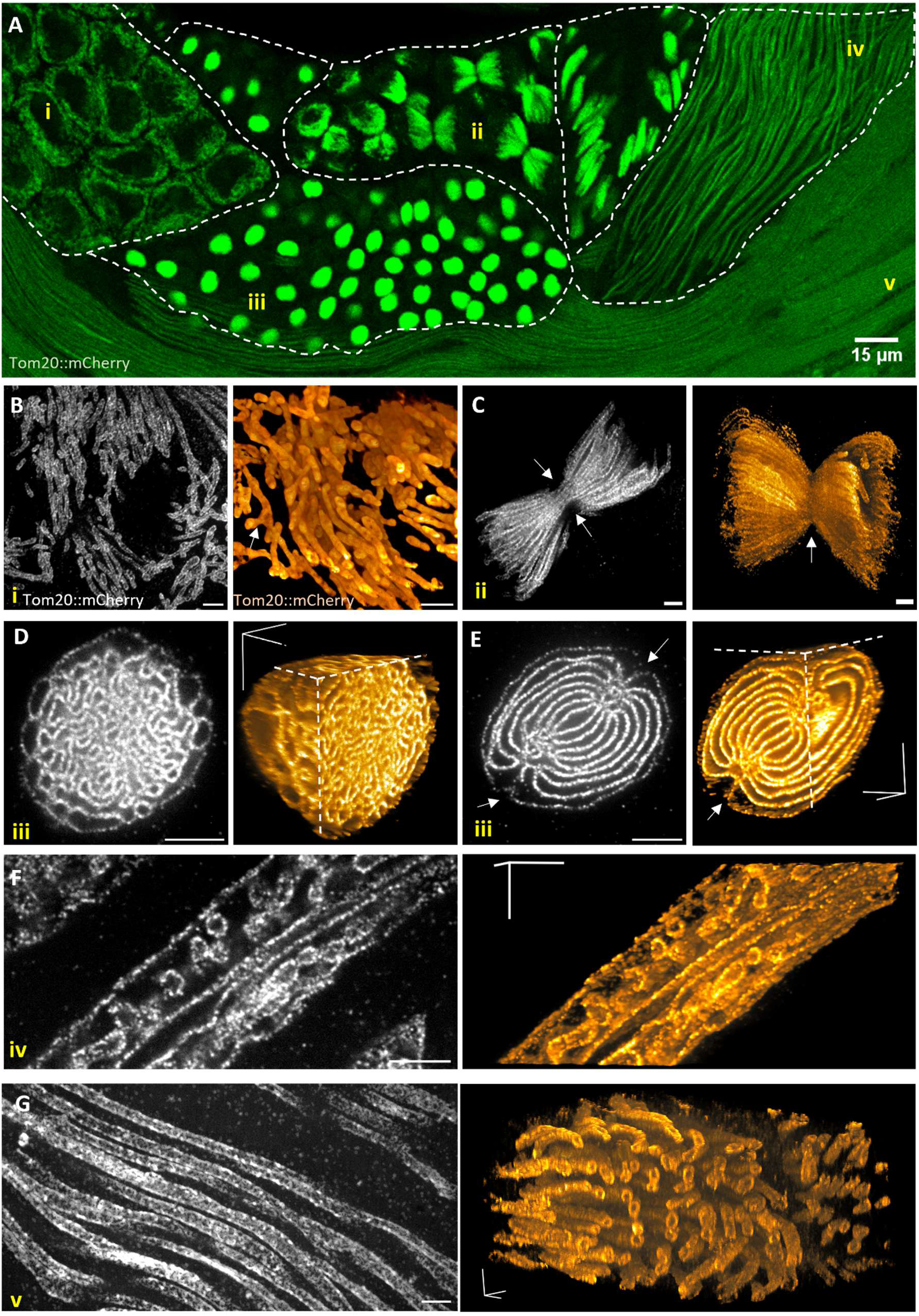
Expansion microscopy reveals finer morphological features of post-meiotic mitochondrial dynamics. (A) Confocal image of adult *Drosophila* testis revealing various developmental stages and their associated mitochondrial morphology. (i) Meiotic stage, (ii) Meiotic division, (iii) Nebenkern, (iv) elongating spermatids, and (v) elongated spermatids. Tom20::mCherry (Green) marks mitochondria. (B-G) Mitochondria imaged using expansion microscopy (Left). Tom20::mCherry (Grey) is used to mark the mitochondria. Volumetric rendering of mitochondrion (Right). (B) Mitochondria in meiotic spermatids. This stage corresponds to region (i) in (A). Arrows indicate branch points. (C) Bow-tie-shaped mitochondria in the dividing meiotic spermatids. This stage corresponds to region (ii) in (A). Arrows indicate constriction along the cytokinetic furrow. (D) “Early nebenkerns” with mitochondria in a spongiform arrangement. This stage corresponds to region (iii) in (A). Volumetric rendering of early nebenkerns clipped along x- and z- planes, displaying a sphere of highly unordered sheet-like mitochondria (Right). (E) “Late nebenkerns” showing ordered whorls of mitochondria with constrictions on either end (Left). Volumetric rendering of late nebenkerns clipped along x- and z- planes, displaying several ordered sheets of mitochondria constricted along a midplane (Right). (F) Divided mitochondrial derivatives in the elongating spermatid corresponding to (iv) in (A) (Left). Volumetric rendering shows vesicular structures within derivatives (Right) (G) Divided mitochondrial derivatives in the elongated spermatid corresponding to (v) in (A) (Left). Volumetric rendering shows the paired arrangement of derivatives (Right) The scale bar represents 5 μm unless otherwise noted.

In the pre-meiotic spermatocytes **(Fig. 1Ai),** mitochondria display short tubular morphologies with occasional branching **(Fig. 1B)**. During meiotic division (**Fig. 1Aii**), tubular mitochondria of similar length aligned symmetrically across the cytokinetic furrow, forming a double-nap cone arrangement **(Fig. 1C)**. Following the second meiotic division, mitochondrial tubules aggregate near one side of the nucleus, marking the onset of nebenkern formation (**Fig. 1Aiii**). Although the nebenkern appears as a homogeneous spherical structure by confocal imaging, expansion microscopy resolved it into two distinct morphologies in the round spermatid stage **(Fig 1D, E)**. One exhibits an unordered, spongiform arrangement of mitochondrial sheets **(Fig 1D)**. In contrast, the second displays a highly ordered organization, with ∼7–8 concentric sheets arranged in a whorl-like (“onion”) configuration, indicating a transition from an initially disordered to a highly ordered lamellar organization during nebenkern maturation **(Fig. 1E)**. This ordered state precedes elongation, during which the nebenkern bifurcates into two mitochondrial derivatives that extend to ∼1.8 mm in length **(Fig. 1A(iv-v), F, G)**.

### Developmental downregulation of outer membrane proteins defines maturation states

To study how mitochondrial remodeling is coordinated during spermiogenesis, we examined the localization of key mitochondrial proteins, including Marf, OPA1, and DRP1 (**Table S1).** We identified two distinct populations of round spermatids, distinguished by levels of four outer mitochondrial membrane (OMM) proteins: Marf::GFP **(Fig 2A)**, Mdi::GFP **(Fig S1A),** Daed::GFP **(Fig S1B)**, and Gasz::GFP **(Fig S1C)**. One population displayed high levels of these proteins, whereas the other showed a marked reduction. These reduced levels persisted during subsequent elongation stages. This transition was consistently observed across multiple Marf reporters, including Marf::mCherry **(Fig S1F)**, Marf::HA **(Fig S1G)**, and anti-Marf staining **(Fig S2H,I)**, confirming that Marf undergoes a developmentally programmed downregulation during nebenkern maturation. Marf levels strongly correlated with Daed::GFP intensity **(Fig. S1D)**, but not with another OMM protein, TOM20::mCherry, which remained constant **(Fig. S1E)**. Together, these results define a sharp developmental transition marked by the coordinated and selective reduction of specific OMM proteins during nebenkern maturation, with Marf downregulation signaling a shift toward a reduced mitochondrial fusogenic state prior to its bifurcation.

**Figure 2.**
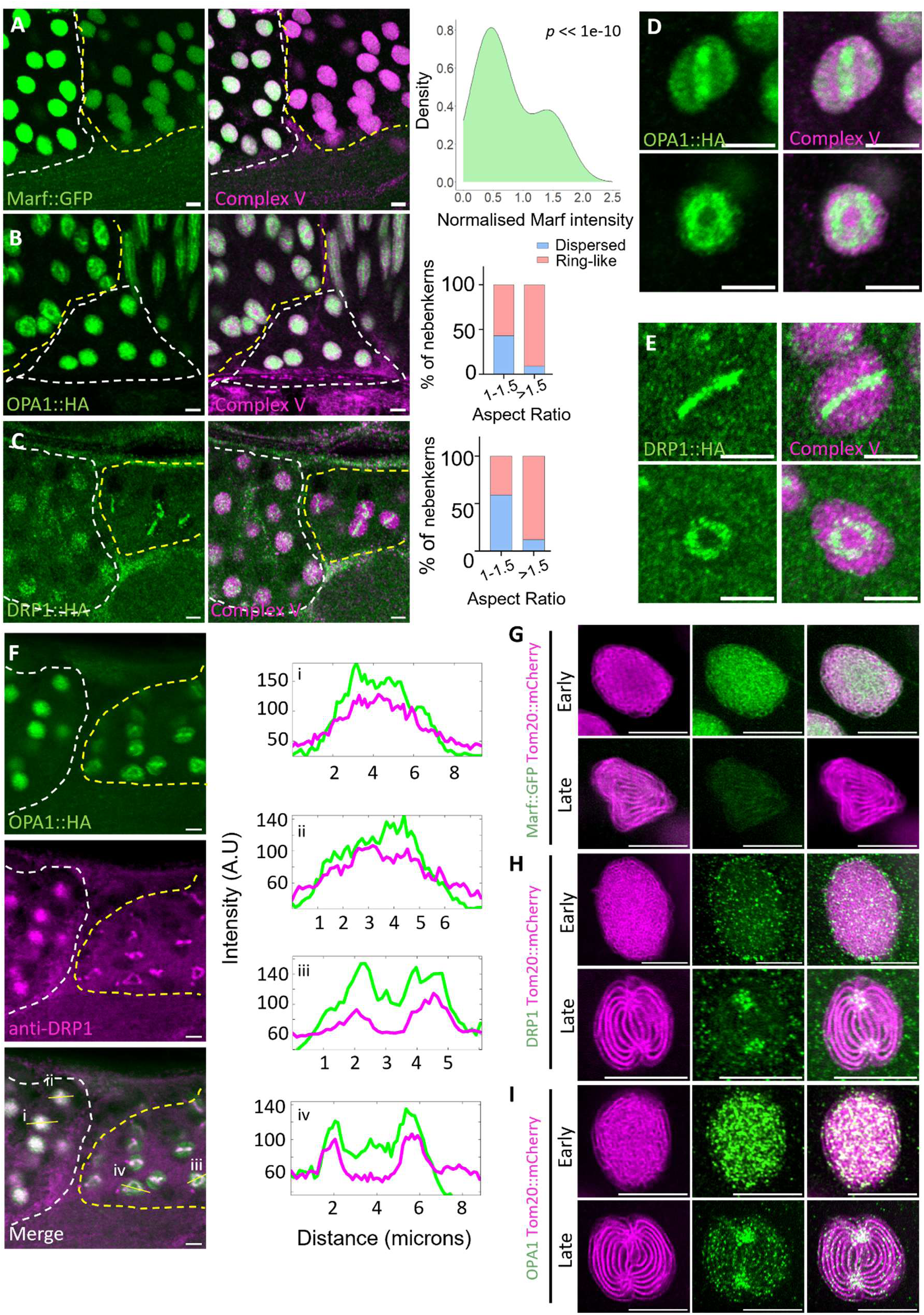
Fission and fusion machinery are coordinately regulated during nebenkern maturation. (A) Two adjacent round spermatid cysts displaying high and low levels of Marf in their nebenkern mitochondria. Density plot of Marf intensity normalized to Complex V intensity (Right). n=184 nebenkerns from 6 testis. P-value suggests a significant deviation from unimodality. Nebenkerns shows Marf::GFP (green) and Complex V (magenta). (B) Two populations of OPA1 showing dispersed and ring morphology. Proportions of nebenkerns with dispersed or ring-like localization of OPA1 are quantified (Right). n=371 nebenkerns from 15 testes. Chi-Squared analysis with Yates’ correction shows a significant association of OPA1 localization change with an increase in the aspect ratio of nebenkerns (p-value: <0.0001). (D) OPA1 rings in two different orientations. OPA1::HA in green, ComplexV in magenta. (C) Two populations of DRP1 showing dispersed and ring morphology. Proportions of nebenkerns with dispersed or ring-like localization of DRP1 are quantified (Right). n=372 nebenkerns from 14 testes. Chi-Squared analysis with Yates’ correction shows a significant association of DRP1 localization change with an increase in the aspect ratio of nebenkerns (p-value: <0.0001). (E) DRP1 rings in two different orientations. DRP1::HA in green, ComplexV in magenta. (F) Colocalization of DRP1 and OPA1 during nebenkern development. Line profiles (correspondingly numbered) measuring OPA1 and DRP1 intensity across nebenkerns. OPA1::HA in green. anti-DRP1 in magenta. (G) Marf levels in early nebenkerns with unordered sheet morphology (Top) and in late nebenkerns with ordered sheet morphology (Bottom). Marf::GFP (green) and Tom20::mCherry (magenta) imaged using SoRa super-resolution microscope with deconvolution. (H) Dispersed localization of DRP1 in early nebenkerns (Top) and enrichment of DRP1 in the constriction sites in late nebenkerns (Bottom). DRP1::HA (green) and Tom20::mCherry (magenta) imaged using SoRa super-resolution microscope with deconvolution. (I) Dispersed localization of OPA1 in early nebenkerns (Top) and enrichment of OPA1 in the constriction sites in late nebenkerns (Bottom). OPA1::HA (green) and Tom20::mCherry (magenta) imaged using SoRa super-resolution microscope with deconvolution. The scale bars indicate 5 μm unless otherwise noted.

### OPA1 and DRP1 assemble into a midplane ring during nebenkern maturation

Interestingly, OPA1 and DRP1, which are required for mitochondrial fusion of the inner mitochondrial membrane (IMM) and fission, respectively, exhibited strikingly similar and dynamic localization patterns in nebenkerns **(Table S1)**.

In early-stage nebenkerns, both proteins appeared as dispersed puncta across the mitochondrial surface. In contrast, in a second population, OPA1 and DRP1 reorganized into a distinct, colocalized midplane-ring encircling the nebenkern **(Fig. 2D,E, Suppl. Video 1,2)**. As nebenkerns progressed toward elongation, the fraction displaying OPA1 and DRP1 rings increased with aspect ratio, indicating that ring formation accompanies maturation **(Fig. 2B,C)**. Moreover, OPA1-ring and DRP1-ring colocalized with each other, suggesting OPA1/DRP1 dynamics are spatially and temporally coordinated during nebenkern development **(Fig. 2F)**

Notably, nebenkerns with dispersed localization of OPA1 and DRP1 exhibited high Marf levels. In contrast, those with ring assemblies exhibited reduced Marf, linking reduced fusogenicity with the emergence of a spatially organized fission assembly **(Fig. S2A,B)**.

To relate these patterns to mitochondrial architecture, we used SoRa super-resolution imaging to distinguish nebenkern morphologies. Unordered, spongiform nebenkern displayed high Marf levels and diffuse OPA1 and DRP1 localization, whereas ordered, lamellar nebenkerns were characterized by reduced Marf and the emergence of a sharply defined OPA1 and DRP1 ring along the midplane **(Fig. 2G-I, S2D,E)**.

Together, these results show that nebenkern maturation is marked by a transition from dispersed to spatially organized OPA1 and DRP1 assemblies, highlighting a transition in which mitochondrial organization possibly defines the spatial positioning of fission machinery.

### Membrane tubulation licenses spatially restricted DRP1 assembly and drives nebenkern division

Late nebenkerns displayed a pronounced constriction along the middle groove **(Fig. 3A)**, coinciding with the localization of OPA1 and DRP1 rings at this site **(Fig. 2H,I)**. To define the structural basis of this constriction, we performed volumetric reconstructions of expanded nebenkerns.

**Figure 3.**
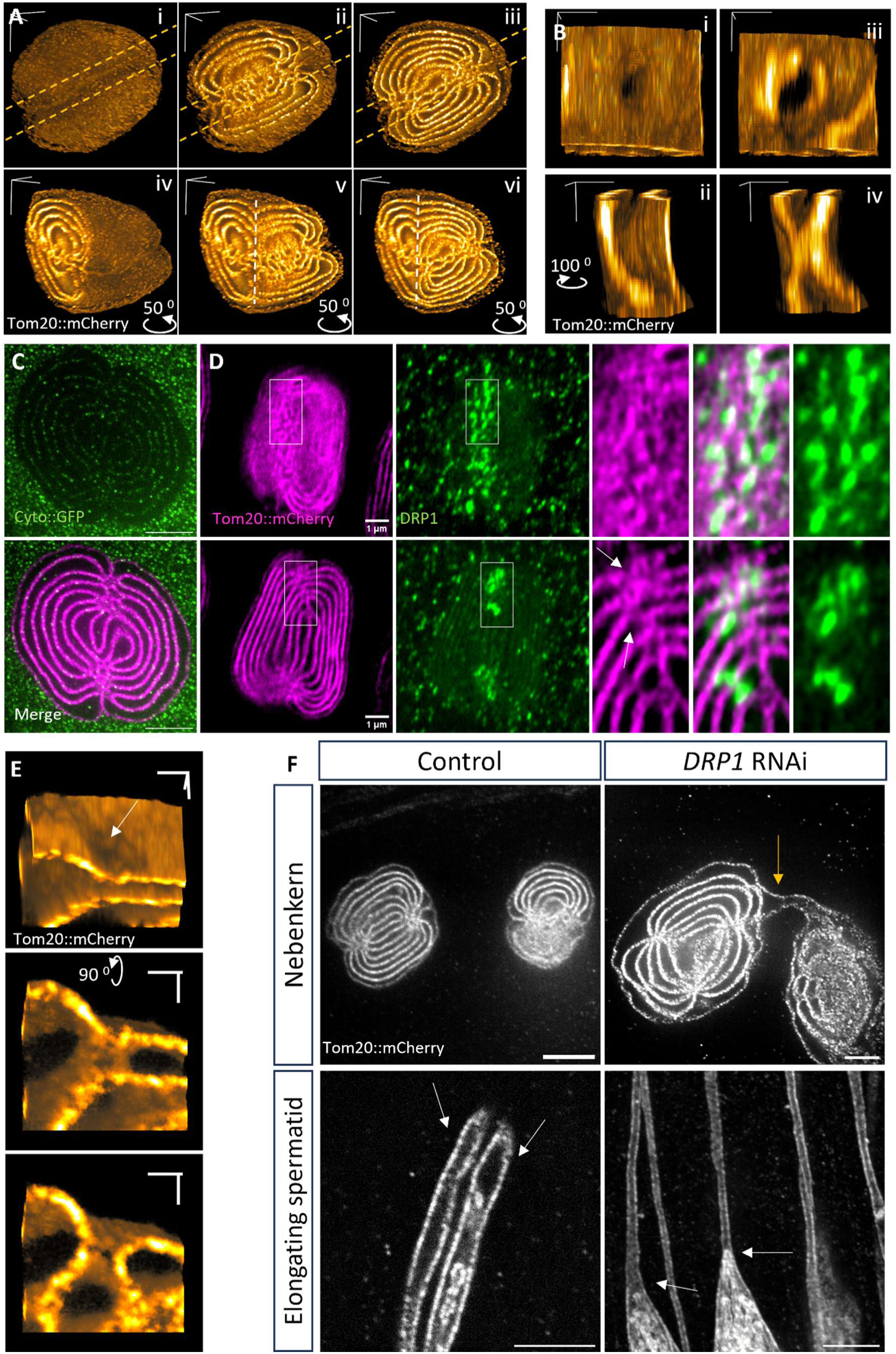
Geometry-licensed DRP1 recruitment regulates nebenkern bifurcation. Tom20::mCherry marks mitochondria. (A) Volumetric rendering of a late nebenkern, marked by Tom20::mCherry. The volumes are unclipped (i) or clipped at different depths along the z-plane (ii and iii), and are rotated 50^0^ along the y-axis and clipped along the x-plane (iv-vi). Constrictions of all sheets are restricted to the middle groove region (marked with orange dashed lines). (B) Volumetric rendering of tubulations of nebenkern sheets resulting from inter-whorl connections in the midplane. The connection resembles a hole from the top view (i) and a tube from the side (ii) (rotated 100° along the z-axis). (iii) Clipping the top sheet in (i) shows a region of positive curvature in the lower sheet. (iv) Clipping (ii) along the x-plane shows the inter-whorl connection constricting the lumen of the middle whorl. (C) A late stage nebenkern after expansion showing cytoplasmic bridges (green - Cytoplasmic GFP) colocalizing with each whorl (magenta - Tom20::mCherry). Scale bar 5ìm. (D) DRP1 localization in the perforated regions of the apical surface of a late nebenkern (Top). DRP1 localization in the medial plane of the same late nebenkern near the constrictions (Bottom). The white box indicates the inset. Arrows indicate constricted regions. DRP1::HA (green) and Tom20::mCherry (magenta). (E) Volumetric rendering of interwhorl connections established in DRP1-knockdown nebenkerns (Bam-GAL4>UAS-DRP1 RNAi). The connections lead to perforations (arrow) on the sheets (Top) and to tubulations (middle and bottom) that lead to constrictions. (F) Nebenkern division (Top) and elongation (Bottom) in DRP1 knockdowns (KD) (Bam-GAL4> UAS-DRP1 RNAi). Nebenkerns divide normally in control flies but remain connected in DRP1-KD flies (yellow arrow). However, membrane ordering and establishment of constrictions proceed normally in DRP1-knockdown nebenkerns. Elongating spermatids in control flies show two mitochondrial derivatives, while in DRP1-KD, a single derivative is observed.

These analyses revealed multiple inter-whorl connections concentrated at the middle groove **(Fig. 3A, Suppl. Video 3)**. Similar connections were occasionally observed elsewhere, but were most prominent at the constriction site. High-resolution rendering of isolated connections showed that they arise from tubular bridges linking adjacent mitochondrial sheets across a central whorl **(Fig. 3B, Suppl. Video 4)**. These bridges generate localized curvature on the inner surface of the sheets and corresponding perforation-like features on the outer surface, indicating membrane tubulation at these sites. These observations suggest that tubulation of mitochondrial sheets through interlocking inter-whorl connections establishes a ring of constrictions around the nebenkern, generating the middle groove **(Fig. S2C)**.

In these renderings, each sheet consists of two sets of OMM-IMM separated by a thin layer of cytoplasm, which could be readily visualized by cytoplasmic GFP **(Fig. 3C, S3A)**. Given that DRP1 preferentially assembles on curved membranes and has not been reported to mediate fission of planar sheets (22, 23), and it could readily access the tubulated constructions in the nebenkern midplane through a thin cytoplasmic layer **(Fig. 3C)**, we hypothesized that these tubulated regions provide a permissive geometry for DRP1 recruitment and stabilization.

Consistent with this, although DRP1 appears as a continuous ring by confocal imaging **(Fig. 2E)**, SoRa super-resolution resolved it into discrete puncta distributed across adjacent whorls within the middle groove **(Fig. 3D, S2D)**. These puncta were enriched at sites displaying perforations in the apical plane and constrictions in the medial plane, features characteristic of tubulated regions identified by expansion microscopy **(Fig. 3D, insets)**. Thus, DRP1 does not form a continuous ring; instead, it preferentially localizes to tubulated membranes within the ring of constrictions around the nebenkern, marking a division plane.

We then tested whether tubulations form before DRP1 localization, and if DRP1 enables nebenkern division. We knocked down DRP1 in the germline cells (Bam-GAL4>UAS-DRP1-RNAi). This caused early mitochondrial segregation defects during meiosis, resulting in interconnected nebenkerns **(Fig. 3F)**. Despite this, nebenkerns still formed ordered sheets and inter-whorl connections with middle-groove constrictions **(Fig. 3E,F)**, indicating that membrane ordering and tubulation occur independently of DRP1.

However, during elongation, these nebenkerns failed to resolve into two distinct mitochondrial derivatives and instead formed a single elongated structure **(Fig. 3F)**. Thus, while DRP1 is dispensable for the formation of constrictions, it is required for the physical separation of the two mitochondrial derivatives. Further, this result implies that an unfurling mechanism, as widely considered (20), would be insufficient to explain nebenkern division. Instead, we propose that nebenkerns actively divide through DRP1-mediated scission along the midplane.

Together, these findings support a model in which membrane ordering generates tubulated geometries that spatially restrict DRP1 assembly at the midplane, highlighting a geometry-dependent mechanism that spatially restricts DRP1 activity to define the plane of mitochondrial division. Given the marked reduction in Marf in these nebenkerns, we next asked whether downregulation of mitochondrial fusogenicity is required to stabilize the division.

### PINK1/Parkin selectively downregulates Marf without affecting nebenkern transitions

While investigating the localization of key mitochondrial proteins **(Table S1)**, we noticed that PINK1 levels increased during nebenkern formation and remained elevated through maturation **(Fig. 4A)**. Stabilization of PINK1 on the outer mitochondrial membrane is known to promote mitophagy, induce Mitofusin degradation, and modulate mitochondrial dynamics in a Parkin-dependent manner (24–27). We therefore asked whether PINK1 regulates nebenkern maturation by controlling Marf levels and/or the spatial organization of OPA1 and DRP1.

**Figure 4.**
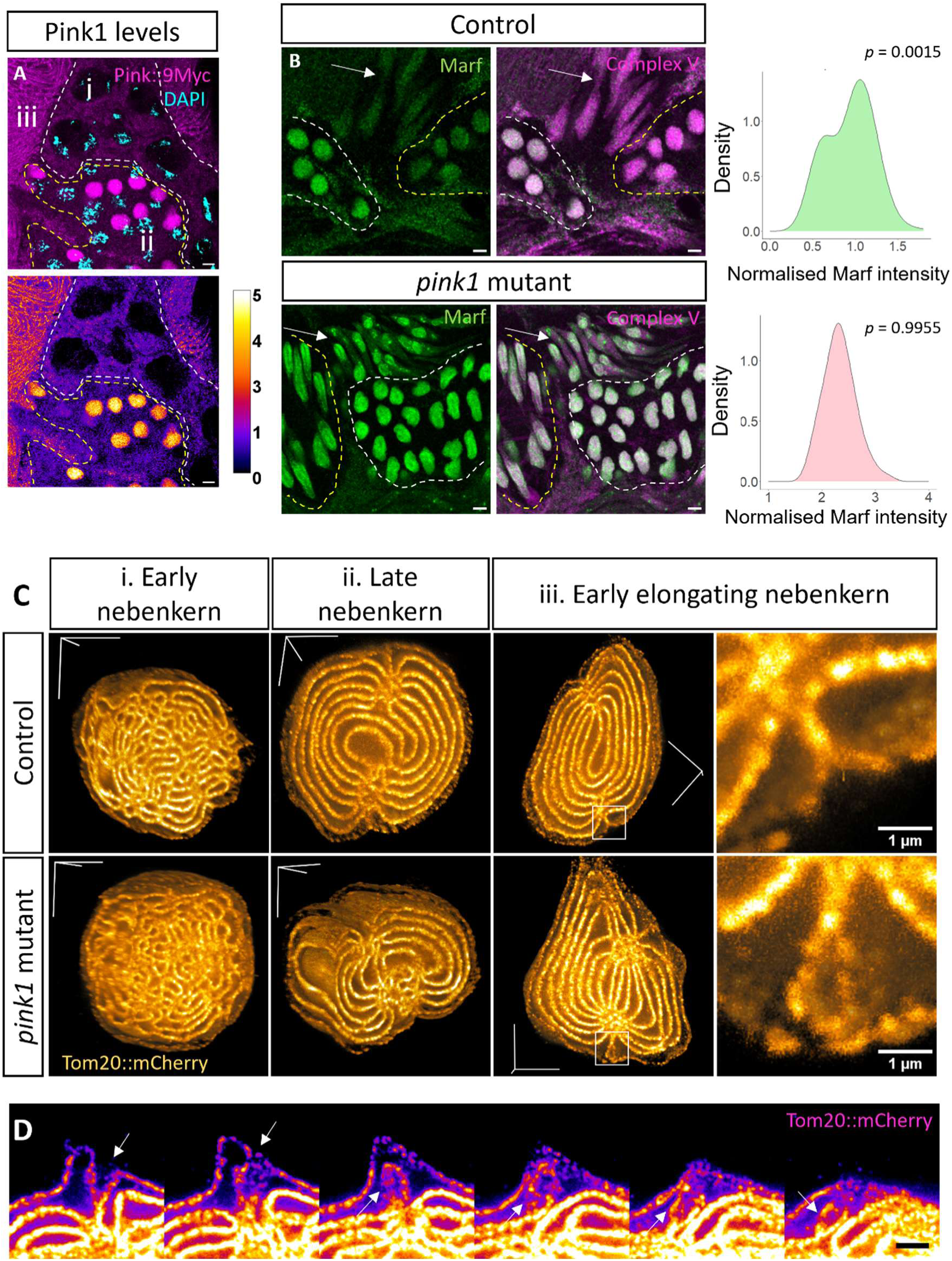
PINK1-mediated Marf downregulation prevents refusion of divided mitochondrial derivatives. (A) PINK1 levels in spermatocytes (i), nebenkerns (ii), and elongating mitochondrial derivatives (iii). PINK1::9Myc is visualized with Myc staining (magenta) and counterstained with DAPI (cyan) (Top). Comparative increase in PINK1 levels during the nebenkern and elongating stages, normalized to PINK1 levels in the spermatocytes and visualized with Fire LUT (Bottom). (B) Marf levels in control (Top) and *pink1* mutants (Bottom). Density profile of Marf normalized to ComplexV in control and *pink1* mutants (Right). n=208 nebenkerns from 6 testis in control and n=197 nebenkerns from 5 testis in *pink1* mutants. Unimodality is estimated by Hartigans’ dip test. P-value <0.05 suggests a significant deviation from unimodality. Marf in green. ComplexV in magenta. Arrows indicate elongating spermatids. (C) Volumetric renderings of early (i), late (ii), and early elongating nebenkern (iii) morphology determined by expansion microscopy and clipped along z- and x-axis in control (Top) and in *pink1* mutants (Bottom). Control nebenkern shows distinct outer whorls, while *pink1* mutants show a fused outer whorl. White box indicates insets. Mitochondrial membranes marked by Tom20::mCherry. (D) Walkthrough along the z-direction of the surface of a *pink1* mutant nebenkern showing looping and re-fusion of the outer whorls. Mitochondrial membranes marked by Tom20::mCherry.

In *pink1* mutants, nebenkerns progressed through ordered morphological transitions **(Fig. 4C)** and formed OPA1/DRP1 rings with normal spatial organization **(Fig. S4C,D)**, indicating that PINK1 is not required for membrane ordering or assembly of the fission machinery.

In contrast, Marf levels remained uniformly high across all stages in *pink1* mutants and failed to distinguish early and late nebenkern populations **(Fig. 4B)**. A similar persistence of elevated Marf was observed in *parkin(park)* mutants **(Fig. S4B)**. Notably, other outer membrane proteins such as Gasz::GFP continued to mark distinct nebenkern populations in *pink1* mutants **(Fig S4A**), indicating that developmental progression occurs despite the absence of Marf downregulation.

Together, these results show that the PINK1/Parkin pathway selectively downregulates Marf during nebenkern maturation without affecting membrane ordering or spatial organization of the fission machinery.

### PINK1-mediated Marf downregulation prevents refusion of dividing mitochondrial derivatives

Loss of PINK1 or Parkin results in male sterility and the formation of a single elongated mitochondrial derivative instead of two, indicating a defect in nebenkern division (25). Given the persistence of high Marf levels in these mutants, we hypothesized that elevated fusogenicity leads to refusion of segregating mitochondrial derivatives.

While early and late nebenkern architecture, including sheet organization and middle-groove constrictions, appeared largely normal in *pink1* mutants **(Fig 4C i,ii)**, defects became apparent during early elongation. In *pink1* mutants, outer whorls frequently appeared fused despite clear constriction and bifurcation of inner whorls **(Fig. 4C-iii; Fig. S5)**, whereas in controls, these whorls remained separated.

Volumetric analysis further revealed tubular extensions from opposing sides of the middle groove that contacted and merged, forming bridges between outer whorls **(Fig. 4D)**, consistent with fusion events between partially segregated mitochondrial derivatives.

PINK1-mediated Marf downregulation during the late nebenkern stage also correlates with the nebenkern-restricted expression of *fuzzy onions* (*fzo*), a paralog of *marf* (*28*), suggesting late nebenkerns have highly downregulated fusogenicity. Together, these observations indicate that in the absence of PINK1-mediated Marf downregulation, elevated fusogenicity permits refusion of dividing mitochondrial derivatives, preventing their stable separation. Thus, while membrane tubulation defines the site of division, PINK1/Parkin-dependent reduction in Marf levels enforces its irreversibility by suppressing fusion.

### Inverted intramitochondrial vesicles mediate membrane redistribution during elongation

During elongation, we observed vesicular structures within mitochondrial derivatives marked by TOM20::mCherry **(Fig. 1F, 5A)**. These structures displayed diverse morphologies, including spherical, tubular, and hybrid tubulo-spherical forms **(Fig. 5B)**, and were detected within the matrix of both inner and outer whorls **(Fig. S6A ii,iii)**, indicating that they can arise throughout the nebenkern.

**Figure 5.**
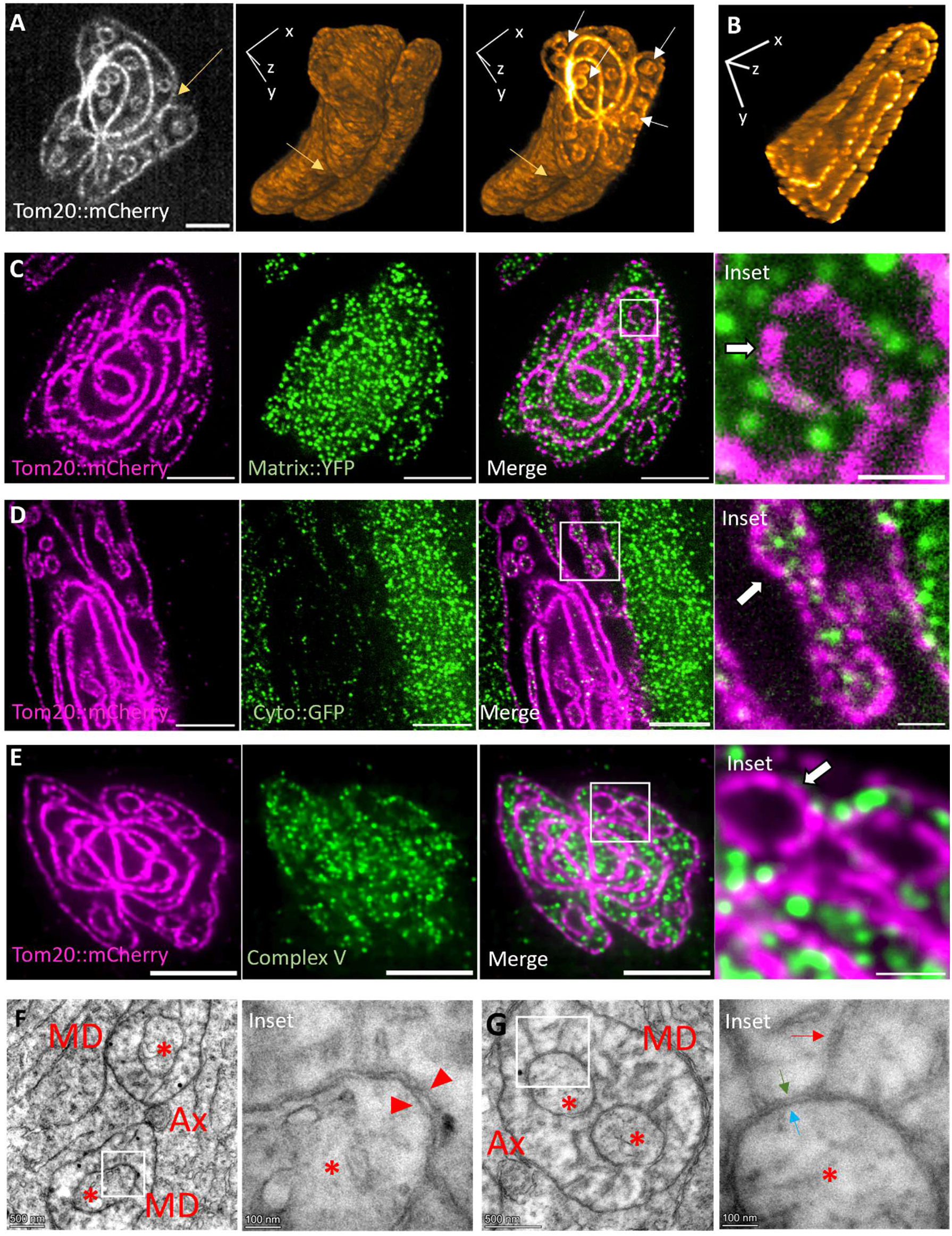
Mitochondria-Derived Inverted Vesicles (MDiVs) form during mitochondrial elongation. (A) Mitochondria in an early elongating spermatid visualized with expansion microscopy (Left). Volumetric rendering of early elongating mitochondria displaying the division furrow (yellow arrow, middle), and clipped along the z-plane displaying vesicle-like structures in the mitochondrial matrix between whorls (white arrows, right). (B) Volumetric rendering of an elongating tip of early spermatid mitochondria displaying tubular and spherical vesicles, clipped along the z-plane. Tom20::mCherry marks mitochondria. (C) Elongating mitochondrial derivatives expressing matrix-targeted YFP (green). OMM is marked with Tom20::mCherry (magenta). White box marks the inset. Arrow indicates MDiV devoid of YFP. (D) Elongating mitochondrial derivatives expressing cytoplasmic GFP (green). OMM is marked with Tom20::mCherry (magenta). White box marks the inset. Arrow indicates MDiV filled with cytoplasmic GFP (green). (E) Elongating mitochondrial derivatives stained with Complex V (green) and Tom20::mCherry (magenta). The lumen of MDiVs (arrows in insets) lacks the Complex V signal. White box marks the inset. (F) TEM image of an elongating spermatid with two well-separated mitochondrial derivatives (MD) and a central axoneme (Ax). MDiVs (asterix) can be observed within both the MDs (Left). The white box marks the inset, which shows the MDiV at higher magnification (Right). The inset shows two membranes forming the MDiV (red arrowheads). (G) TEM image of an elongating spermatid with an axoneme (Ax) and mitochondrial derivative (MD) containing two MDiVs (asterix) visible. The white box marks the inset, which shows the MDiV at higher magnification (Right). Red arrows mark cristae emerging from the outer surface of the MDiV, indicating the inverted nature of MDiVs, with IMM (green arrow) on the outside and OMM (blue arrow) on the inside.

To determine their membrane organization, we leveraged the differential fluorescence intensity of TOM20::mCherry, which labels the outer mitochondrial membrane (OMM). Regions containing closely opposed double membranes, such as inner whorls, exhibited higher signal intensity than single-membrane regions such as the outer surface **(Fig. S6A i, ii, iii)**. Notably, the vesicles displayed fluorescence intensity comparable to single OMM regions, suggesting that they are bound by a single TOM20-positive membrane **(Fig. S6A ii, iii)**. Transmission electron microscopy revealed that these vesicles are enclosed by double membranes **(Fig. 5F)**, indicating the presence of both OMM and IMM. However, they lacked mitochondrial matrix, as evidenced by the absence of matrix-targeted YFP **(Fig. 5C)**.

These observations suggested that the vesicles might form by inward budding of mitochondrial membranes, generating an inverted topology. To test this, we performed expansion microscopy in flies expressing cytoplasmic GFP alongside TOM20::mCherry. GFP signal was detected within the vesicular lumen but not within mitochondrial sheets, indicating that these vesicles enclose cytoplasmic material **(Fig. 5D)**. Consistently, IMM marker (Complex V) was absent from the vesicle lumen **(Fig. 5E)**. Electron microscopy further revealed cristae-like structures oriented outward from the vesicle surface **(Fig. 5G)**, consistent with inversion of membrane topology.

Together, these results demonstrate that these vesicles possess an inverted architecture, with the IMM facing outward and the OMM enclosing cytoplasmic content **(Fig. 6F)**. We therefore term them Mitochondria-Derived Inverted Vesicles (MDiVs), representing a previously unrecognized class of intramitochondrial vesicles distinct from canonical mitochondria-derived vesicles (MDVs) **(Suppl. Video 5)**.

**Figure 6.**
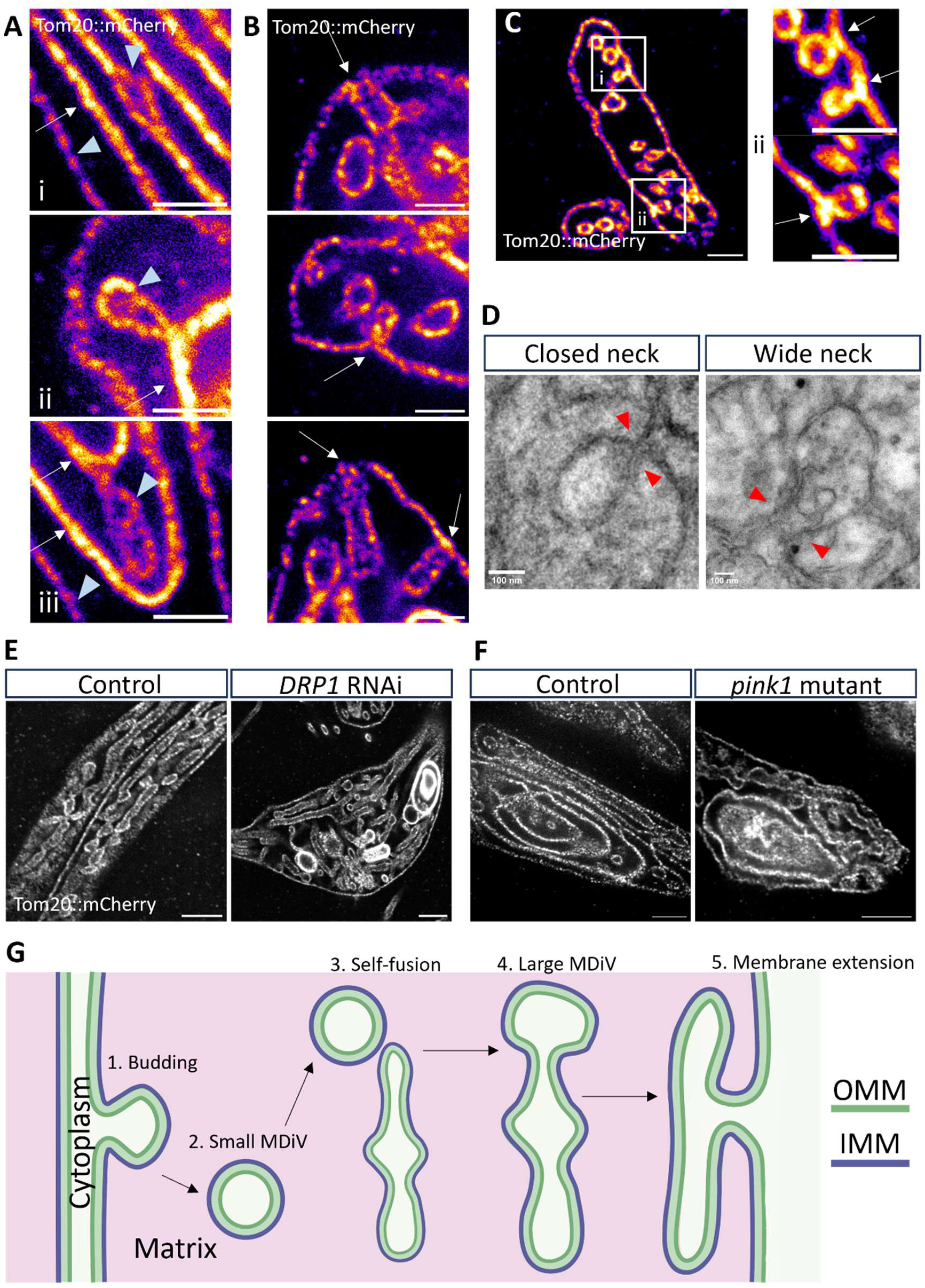
MDiVs may mediate membrane redistribution during mitochondrial elongation. (A) Budding MDiVs. Mitochondria are marked with Tom20::mCherry and visualized with Fire LUT. Bulging between two adjacent whorls during the initial stages of MDiV formation (i). Budding MDiVs are attached to the whorls at different stages of formation (ii and iii). High intensity regions (white arrows) suggest proximity of two OMMs, while low intensity regions (arrowheads) suggest a single OMM. (B) Examples of MDiVs in close proximity to the extending outer whorls. Arrows indicate points of contact. (C) Potential simultaneous fusion events of multiple MDiVs inside an MD. Arrows indicate points of contact. White boxes correspond to insets i and ii. Scale bars 2 μm (A-C). (D) TEM images of MDiVs in elongating spermatids at different stages of contact. A closed-neck conformation (Left) and a wide-neck conformation (right). Red arrowheads point to the region of contact between MDiV and the outer whorl. (E) Expansion microscopy images of elongating mitochondrial derivatives with MDiVs suspended within. MDiVs can be observed in both the control (Bam-GAL4) and the DRP1 knockdown (BamGal4>UAS-DRP1 RNAi) conditions. (F) Expansion microscopy images of elongating mitochondrial derivatives with MDiVs suspended within. MDiVs can be observed in both the control and *pink1* mutants. Scale bars are 5μm (E-F). Tom20::mCherry marks mitochondria. (G) Schematic of the proposed model of MDiV-mediated membrane redistribution. MDiVs form through inverted budding events (1). They are suspended within the mitochondrial matrix and possess an inverted membrane topology, and have trapped cytoplasm (2). Free-floating MDiVs may fuse to each other, giving rise to tubulo-spherical morphology (3), causing an overall increase in their size (4). These large MDiVs are reintegrated with the mitochondrial membrane through IMM-IMM fusion, leading to membrane redistribution and extension (5).

MDiVs emerged specifically during elongation, with their abundance increasing as the number of mitochondrial whorls decreased **(Fig. S6A)**, suggesting a link to membrane remodeling. Consistent with this, we observed inward budding events from mitochondrial sheets resembling endocytic-like invaginations **(Fig. 6A)**, indicating that MDiVs form through active membrane remodeling.

Notably, MDiVs were absent in fully elongated mitochondrial derivatives **(Fig. S6A vi)**. During elongation, they were frequently observed in proximity to the outer whorl, with instances of apparent fusion between MDiVs and the outer membrane **(Fig. 6B,C, Suppl. Video S6)**. Electron microscopy further supported membrane continuity between MDiVs and the outer whorl **(Fig. 6D).**

These observations suggest that MDiVs may mediate membrane transfer from inner whorls to the elongating outer mitochondrial derivative. While the static nature of these measurements precludes direct inference of directionality, the spatial and temporal dynamics of MDiVs are consistent with a vesicle-mediated mechanism of intramitochondrial membrane redistribution **(Fig 6F).**

Finally, MDiV formation persisted in DRP1-depleted and PINK1-deficient spermatids **(Fig. 6E,F)**, indicating that, unlike MDV formation (29, 30), this process is independent of canonical fission machinery and the PINK1 pathway. We suggest that MDiVs may emerge through tubulation and budding, akin to MDVs, presenting similarities in membrane remodeling during their emergence (31). However, membrane remodeling machineries regulating MDV formation may differ from MDiV formation. Based on the mechanism of MDiV emergence, we speculate that matrix-localized factors may drive MDiV formation.

## Discussion

Mitochondria undergo extensive morphological remodeling during development, yet how they are reorganized in a coordinated and spatially controlled manner remains poorly understood. Using *Drosophila* spermatogenesis as a model, we identify a program that enables the division and elongation of the nebenkern, a giant mitochondrial structure formed by the fusion of all mitochondria in the cell. Our findings reveal that this process is achieved through the coordinated action of three mechanistic modules: geometry-dependent positioning of the fission machinery, regulated suppression of membrane fusion, and vesicle-mediated membrane redistribution.

### Spatial patterning of mitochondrial division

A central challenge in remodeling the nebenkern arises from its multilamellar architecture, in which multiple concentric mitochondrial sheets must be partitioned in a coordinated manner. We find that membrane ordering generates interlinked sheets that undergo localized tubulation at the midplane, creating regions of high curvature **(Fig. 3A,B)**. These tubulated geometries coincide with the assembly of DRP1 into a ring-like structure **(Fig. 3D)**, suggesting that membrane curvature licenses spatially restricted recruitment and stabilization of the fission machinery. In this context, DRP1 does not initiate membrane tubulation but instead acts downstream of geometry, executing division at pre-patterned sites **(Fig. 3E,F)**. This is consistent with previous studies showing that membrane constriction to diameters of ∼10–250 nm facilitates higher-order DRP1 oligomerization and membrane scission (22). This supports a model in which mitochondrial architecture itself encodes spatial information that defines the plane of division. Such changes in local geometry may prove critical during the division of large mitochondrial structures.

### Fine-tuning fusogenicity

While DRP1-mediated constriction defines where division occurs, successful separation of mitochondrial derivatives requires suppression of opposing fusion forces. We show that the PINK1/Parkin pathway downregulates the Marf/Mitofusin during nebenkern maturation **(Fig. 4B)**, without affecting membrane ordering **(Fig. 4C i,ii)** or DRP1 ring formation **(Fig. S4D)**. In the absence of PINK1 or Parkin, Marf levels remain elevated, and although constriction proceeds, the two mitochondrial derivatives fail to remain separated and instead undergo refusion **(Fig. 4Ciii, D, S5).** These findings indicate that PINK1-dependent reduction in fusogenicity is not required to initiate division but is essential to stabilize it, enforcing the irreversibility of the process. This uncoupling of spatially patterned fission and the suppression of fusion reveals that mitochondrial division is controlled through parallel but functionally distinct pathways.

Although PINK1 is best known for its role in mitophagy (32), studies from our laboratory have suggested that PINK1-mediated downregulation of Marf/Mitofusin also promotes the segregation of defective mitochondria to preserve cellular homeostasis (33). Together, these findings suggest that regulation of mitochondrial fusogenicity may represent a central and evolutionarily conserved function of the PINK1/Parkin pathway, operating across both developmental and stress-induced remodeling contexts.

### Membrane remodeling during mitochondrial elongation

Following division, elongation of the mitochondrial derivatives presents an additional challenge, requiring redistribution of membrane from a multilamellar structure into extended tubular forms. This process appears largely independent of canonical Marf- and DRP1-mediated fusion-fission dynamics, as Marf is downregulated in these stages **(Fig. 2A)**, and resolution and elongation proceed in DRP1-knockdown **(Fig. S6B)**. We identify MDiVs as a previously unrecognized intermediate in this remodeling process. These vesicles arise through inward budding of mitochondrial membranes **(Fig. 6A)**, exhibit inverted topology **(Fig. 5F,G)**, and transiently accumulate during elongation **(Fig. S6A)**. Their absence in fully elongated spermatids suggests that they are transient intermediates that are ultimately incorporated into the mitochondrial derivatives. Although the static nature of our observations precludes direct visualization of MDiV dynamics or membrane transfer, their spatial and temporal association with elongating mitochondrial derivatives supports a role in membrane redistribution during morphogenesis **(Fig 6G)**. The molecular mechanisms governing MDiV biogenesis and fusion remain important questions for future investigation.

Although not previously characterized in detail, MDiV-like intramitochondrial membrane-bound structures have been reported in certain contexts. For example, a recent study of stress-induced megamitochondria described lysosomal engulfment through deep mitochondrial membrane invaginations (34). Similarly, loss of *eat-3* (the *C. elegans OPA1* ortholog) results in the accumulation of double-membrane compartments within mitochondria (35). Furthermore, enlarged mitochondria in the muscles of aging *Drosophila* have also been reported to contain cytoplasmic inclusions bound by double membranes (36). While the topology and origin of these structures are not well understood, these observations suggest that MDiV-like membrane remodeling may represent a broader mitochondrial phenomenon operating across physiological and pathological contexts. Our findings provide a framework for reinterpreting these previously observed intramitochondrial membrane-bound structures, which may form via inward membrane budding.

While our data suggest that MDiVs participate in mitochondrial remodeling during spermatid elongation, whether similar structures mediate membrane redistribution, cargo transfer, or quality-control functions in other systems remains an important question for future investigation.

### Developmental coordination of mitochondrial remodeling

Our findings support a model in which mitochondrial remodeling during spermiogenesis emerges from the coordinated action of three mechanistically distinct processes. Membrane geometry establishes spatial cues that position the fission machinery, regulated suppression of fusion ensures that division proceeds unidirectionally, and vesicle-mediated remodeling facilitates the redistribution of membrane required for subsequent elongation. Together, these processes overcome the geometric and stoichiometric constraints inherent to remodeling a giant organelle.

Interestingly, these modules are activated and suppressed during distinct developmental windows; however, synchronously across all 64 spermatids in a given cyst. Moreover, each module works independently of the others, with the success of one module having no bearing on the others. Such a high degree of developmental coordination, yet independent regulation, suggests the existence of higher-order developmental coordination mechanisms, potentially mediated through non-cell-autonomous signals from surrounding somatic cyst cells.

## Materials and Methods

### Fly husbandry

Flies were cultured on standard media containing sucrose, malt, yeast and corn flour at room temperature. Crosses were maintained at 25°C. Crosses involving RNAi were maintained at 28°C. The genotypes used in this study are provided in Table 1.

**Table 1:**
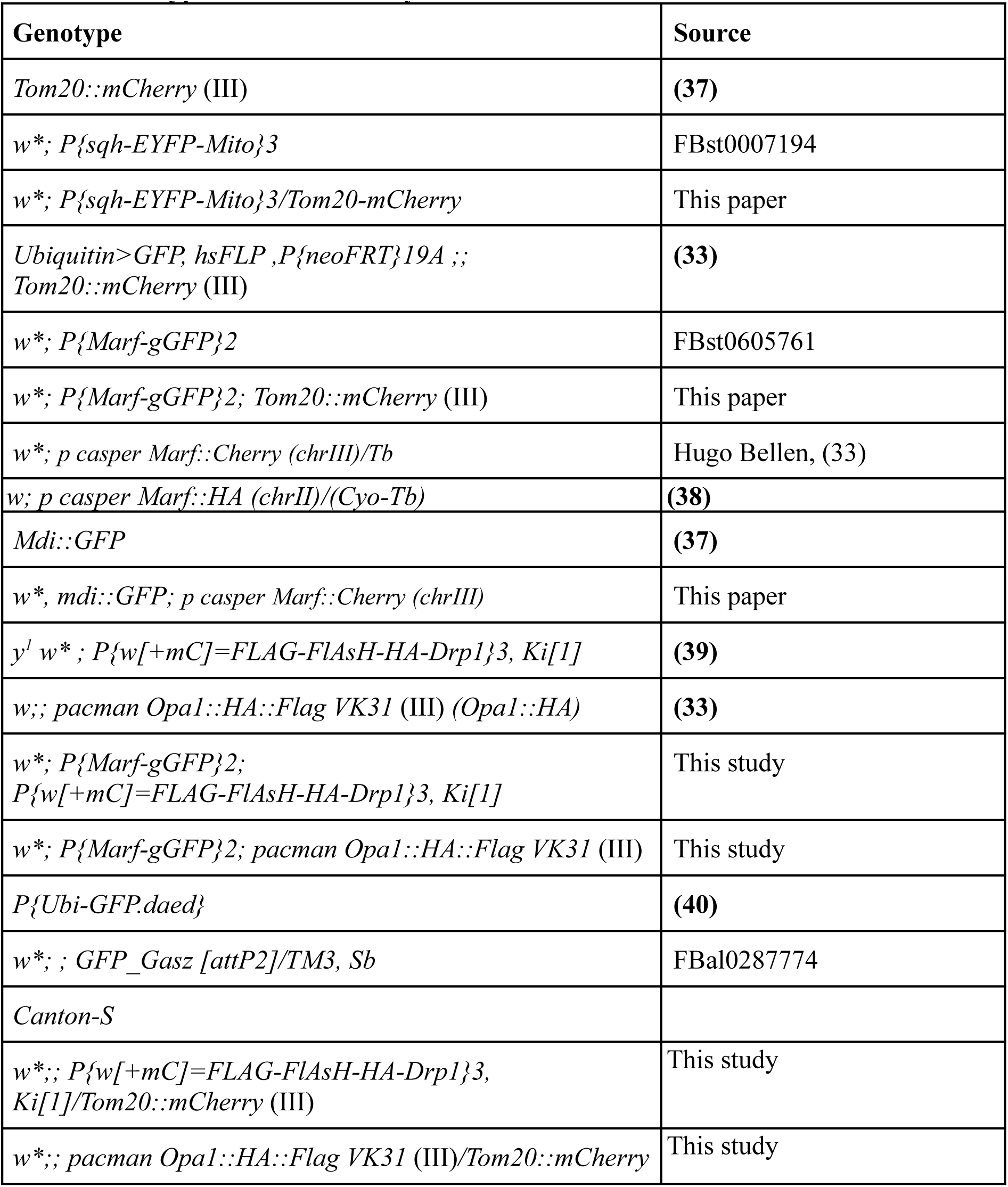
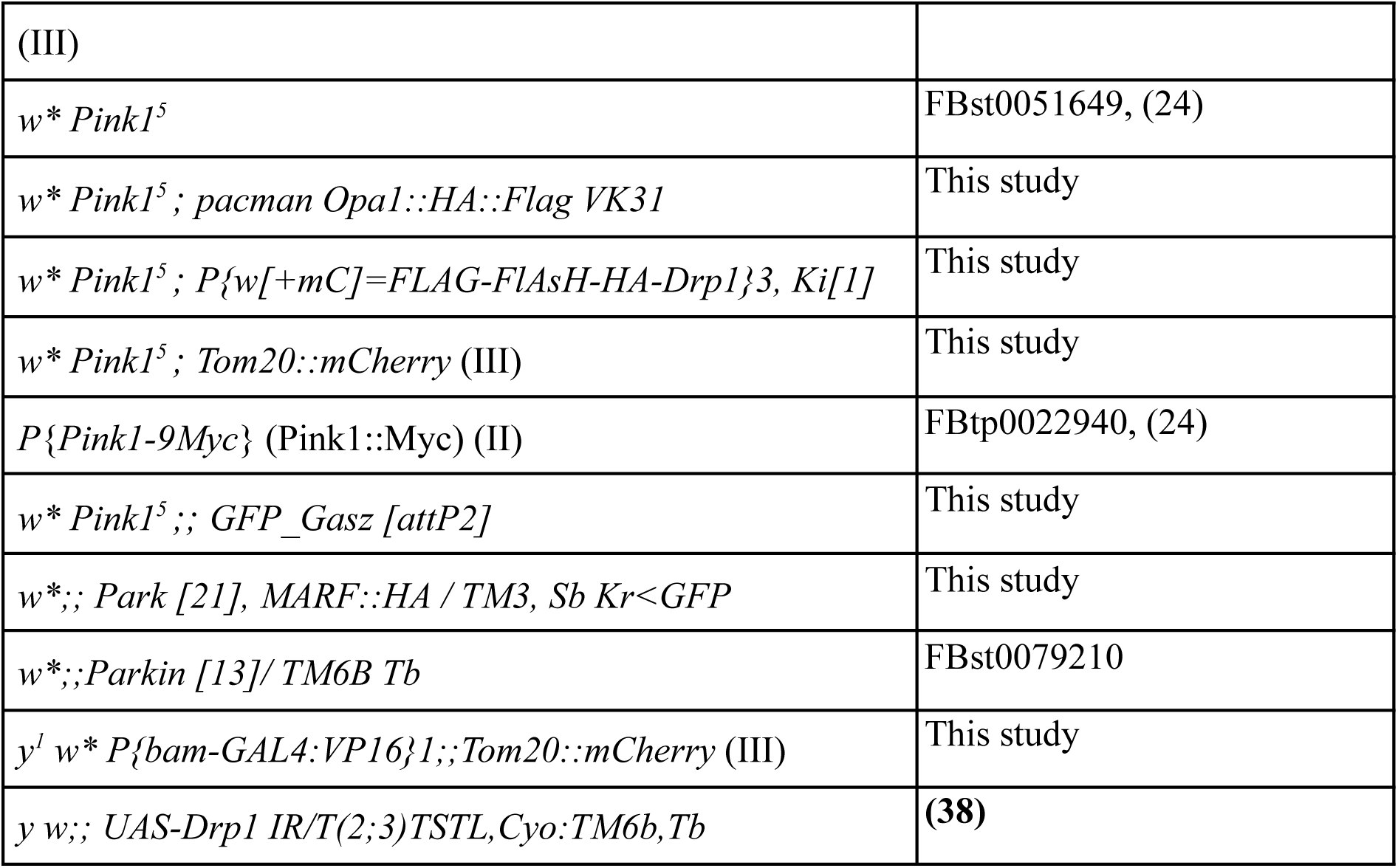
Genotypes used in the study.

### Expansion microscopy of *Drosophila* testis

The expansion microscopy protocol was adapted from (41). Briefly, immunostained testis were incubated in 0.1 mg/ml Acryloyl-X SE (AcX) (Invitrogen A20770) in PBS (prepared fresh from 10 mg/ml AcX in DMSO (Sigma 276855) at room temperature overnight. Following this, the tissues were washed in 1X PBS twice for 15 minutes each. After washing, the tissues were incubated in the gelling solution (Stock X (Table 2), 0.5% 4HT (Sigma 176141), 10% TEMED (Himedia MB026), 10% APS (Himedia MB003), in 47:1:1:1 ratio) at 4°C for 30 minutes in the dark. For polymerization, the tissues were transferred to slides containing 200-300 μl of gelling solution and gently compressed using a parafilm-wrapped coverslip. The gelation was performed by incubating this setup at 37°C for two hours in a humidified chamber. After polymerization, the gel was cut into smaller pieces containing not more than two tissues per piece, and incubated overnight in the digestion buffer (Table 3) containing Proteinase K (Himedia MB086) (final concentration of 8U/ml) for digestion. The digested gel pieces were then stored in 1X PBS at 4°C prior to expansion. Expansion of the digested gel was performed just before imaging. For expansion, the gels were incubated in milliQ water in a large petri dish. Water changes were performed at an interval of 20 minutes thrice. The expanded gels were carefully transferred to a cover glass, and extra liquid was removed with tissue paper. To prevent the gel from drifting while imaging, 0.5% agarose solution (Sigma-Aldrich A9045) was added to the boundaries of the gel.

**Table 2:**
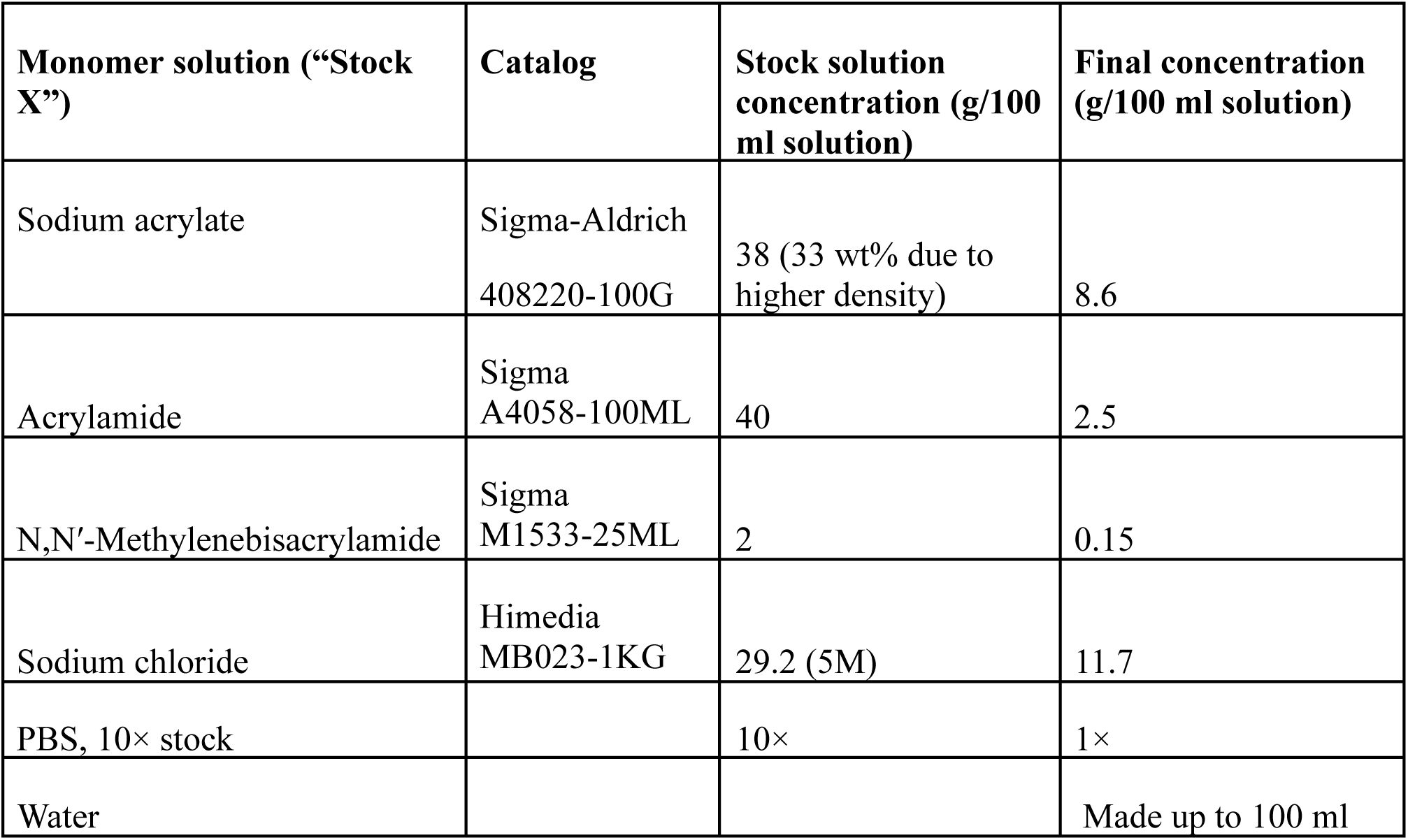
Composition of Stock-X solution.

| **Monomer solution (“Stock X”)** | **Catalog** | **Stock solution concentration (g/100 ml solution)** | **Final concentration (g/100 ml solution)** |
| --- | --- | --- | --- |
| Sodium acrylate | Sigma-Aldrich<br>408220-100G | 38 (33 wt% due to higher density) | 8.6 |
| Acrylamide | Sigma<br>A4058-100ML | 40 | 2.5 |
| N,N'-Methylenebisacrylamide | Sigma<br>M1533-25ML | 2 | 0.15 |
| Sodium chloride | Himedia<br>MB023-1KG | 29.2 (5M) | 11.7 |
| PBS, 10× stock |  | 10× | 1× |
| Water |  |  | Made up to 100 ml |

**Table 3:**
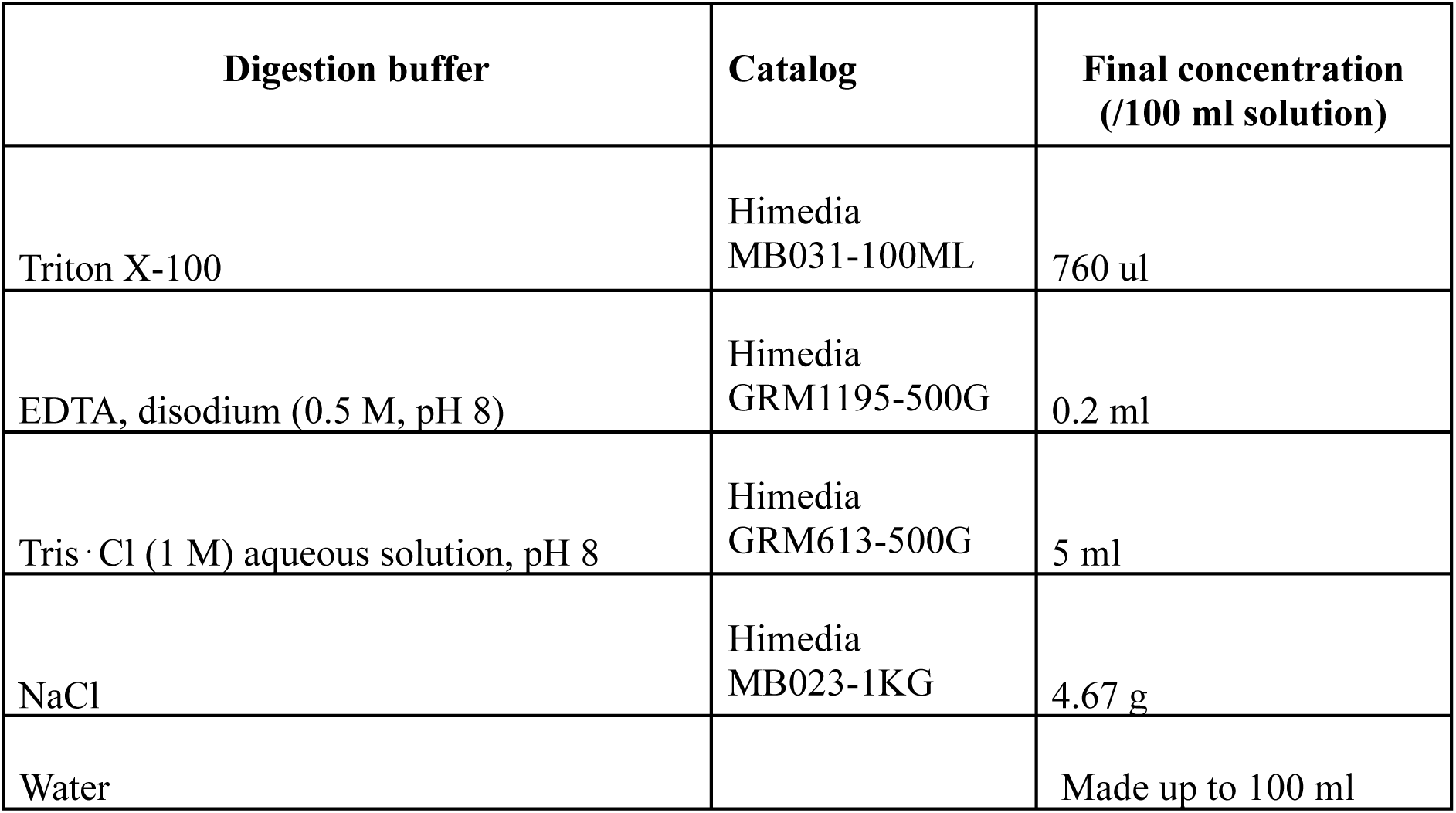
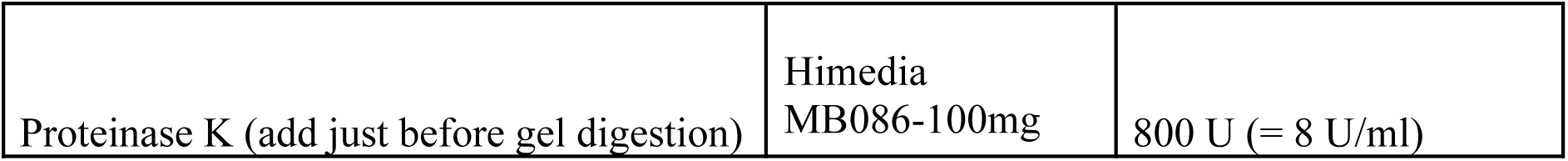
Composition of Digestion buffer.

Imaging of the expanded gels was performed using a 60X water immersion objective with 4X magnification by Lens-switched light path in SORA (Spinning Disk Super Resolution by Optical Pixel Reassignment) mode using an inverted Nikon CSU-W1 SORA spinning disk confocal microscope. A few drops of milliQ were added frequently to moisturize the gel and prevent it from shrinking during the imaging.

### Immunostaining

Testis were dissected in 1X PBS and fixed with 4% paraformaldehyde solution in 1X PBS (Himedia TCL119). Following this, the testis were washed with 0.2% PBST (Triton X-100 (Himedia MB031) in 1X PBS) and incubated with primary antibodies of indicated dilutions overnight at 4°C. This was followed by three washes in 0.2% PBST. The testes were incubated in blocking solution containing 5% Goat serum diluted in 0.2% PBST for 2 hours at room temperature. Following blocking, the testes were incubated in secondary antibodies of the indicated dilutions for 2 hours (overnight at 4^0^ C for expansion microscopy samples) at room temperature. The testes were then washed with 0.2% PBST thrice at room temperature, ten minutes each. The testes were then mounted on a glass slide with a drop of VectaShield (Vector labs H-1900-10). All antibody dilutions were made in 0.2% PBST. A list of antibodies used in the study is provided in Table 4.

**TABLE 4.**

| Antibody | Catalog | Dilution |
| --- | --- | --- |
| Mouse HA | Cell Signaling Technology- 2367S | 1:500 |
| Rabbit HA | Cell Signaling Technology- 3724S | 1:500 |
| Mouse ATP5A | Abcam- ab14748 | 1:500 |
| Rabbit mCherry | Cell Signaling Technology- 43590S | 1:500 |
| Mouse Myc Tag | Cell Signaling Technology-2276S | 1:1000 |
| Rabbit Anti-GFP | Invitrogen- A6455 | 1:500 |
| Chicken Anti-GFP | Abcam- ab13970 | 1:500 |
| GFP Booster | Chromotek- gb2AF488-10 | 1:250 |
| RFP Booster | Chromotek- rb2AF568 | 1:250 |
| Rabbit Marf | This study | 1:2000 |
| Anti-Mouse 488 | Invitrogen- A32723 | 1:500 |
| Anti Mouse 555 | Invitrogen- A21424 | 1:500 |
| Anti-Mouse 633 | Invitrogen- A21052 | 1:500 |
| Anti- Mouse 647 | Invitrogen- A32728 | 1:500 |
| Anti-Rabbit 488 | Invitrogen- A32731 | 1:500 |
| Anti-Rabbit 555 | Invitrogen- A32732 | 1:500 |
| Anti-Rabbit 647 | Invitrogen- A32733 | 1:500 |
| Anti- Chicken 488 | Invitrogen- A32931 | 1:500 |
| Rabbit anti-DRP1 | Gift from Prof. Richa Ricky | 1:500 |
| Rabbit anti-Hsp60 | Cell Signaling Technology- 4870S | 1:500 |

### Image processing and analysis

#### 3-D rendering

Volumetric rendering of all images was performed using “3D script” plugin in ImageJ (42). To increase the smoothness of the rendered volumes along the z-direction, images were scaled along the z-direction with a factor of 5.0 using Bilinear Interpolation without altering the aspect ratio. Background in the projected images was removed by marking the regions using the “free-hand” tool in ImageJ, followed by the “clear selection” dialog in 3D script. For rendering with multiple channels, the “Combined transparency” function was used for visualization.

#### Deconvolution

Images were deconvolved in NIS-Elements software (5.42.01 (Build 1793)) using the Richardson-Lucy algorithm. The number of iterations was determined automatically.

#### Segmentation of nebenkerns

Segmentation of nebenkerns was performed in ImageJ using a custom macro. The nebenkern-containing regions were marked with the “free-hand tool” in the Z-projected images (maximum projection). The region was then duplicated, and the unmarked areas were removed using the “Clear Outside” tool. We performed Gaussian Blur (sigma = 5) followed by auto-thresholding to create masks of the nebenkerns. The masks were filtered using the size parameter set to 10-Infinity to identify the nebenkerns. The masks were overlaid on the original z-projected images to measure Mean intensity and Aspect Ratio across all channels using the “multi-measure” tool.

#### Identification of distinct nebenkern populations

To determine if the observed normalized intensity values of nebenkerns were sampled from different populations, we plotted these values as a probability density function using R. The bandwidth was estimated using the Freedman-Diaconis rule. We then performed Hartigans’ dip test to determine if the curve was unimodal or multimodal. A multimodal curve would indicate the presence of distinct populations of nebenkerns.

### Electron microscopy

Testis were dissected and fixed in Karnovsky’s fixative - 2% Paraformaldehyde (EMS - 15710), 2.5% glutaraldehyde (EMS - 16320) in 0.1M sodium cacodylate buffer (EMS- 12300, pH - 7.4), overnight at 4℃. Fixed samples were washed 5 times with 0.1M cacodylate buffer, 15 minutes per wash. Secondary fixation was done in 1% OsO4 (EMS - 19150) and 1% potassium ferrocyanide for 1h on ice. This was followed by 5 washes with 0.1M cacodylate buffer, each wash lasting 15 minutes. After washing, the samples were dehydrated in graded ethanol and then infiltrated with Spurr resin (EMS - 14300). Samples were embedded and polymerised overnight at 70℃. Ultrathin sections (70nm) were cut using a Leica EM UC7 Ultramicrotome and mounted on copper grids (EMS- 300 Mesh). Imaging was done using Thermo Fisher Talos L120C TEM (120kV).

### Statistical analysis

In all experiments, *n* represents the number of nebenkerns. The number of testis used for each experiment is mentioned in the figure legends. Deviation from unimodality in all density plots was determined using Hartigan’s dip test in R. P-values below 0.05 indicate a significant deviation from unimodality. Categorical data were measured for association using the Chi-Squared test with Yates correction. P-values below 0.05 indicate a significant association. All other data were analyzed using two-tailed unpaired *t*-tests with Welchs’ correction. Chi-Squared analysis and *t*-tests were performed using GraphPad Prism 10.

## Acknowledgements

We thank Hong Xu, Ming Guo, Hugo J. Bellen, Richa Rikhy, and Gregory Hannon for generously sharing fly lines and antibodies. We thank Padmasini Chary and Rajeshwari Krishnamurthy (Bioklone Biotech Private Limited) for assistance with Marf antibody generation. Fly lines obtained from the Vienna Drosophila Resource Center and the Bloomington Drosophila Stock Center (NIH P40OD018537) were used in this study. We also acknowledge the use of FlyBase (flybase.org), a comprehensive database for Drosophila genetics and genomics, which was instrumental in the design and interpretation of this study. We thank the CSIR-CCMB Electron Microscopy Centre for help with electron microscopy. The MJ laboratory is supported by the Department of Atomic Energy, Government of India (Project Identification No. RTI 4007), the Department of Science and Technology, SERB/ANRF (CRG/2020/003275 and CRG/2023/005377), and the Department of Biotechnology (BT/PR32873/BRB/10/1850/2020). A.H. was supported by the Indo-German Grant by the Department of Biotechnology IC-12025(11)/2/2020-ICD-DBT. We thank Anand Vaidya, Tamal Das, Rajit Narayanan, Tarana Anand, Sunayana Sarkar and SK Yasir Hosen for valuable discussions and comments on the manuscript.

## Author Contribution

A.H. and M.J. conceived the study. A.H. designed and performed experiments, analyzed data, and wrote the manuscript. A.H. and V.K. conducted the expansion microscopy experiments. V.M. performed Electron Microscopy. M.J. supervised the project, secured funding, provided overall guidance, and contributed to manuscript writing and revision. All authors discussed the results and approved the final version of the manuscript.

