## Supplemental figures for "Coordinated membrane remodeling and fission-fusion drive mitochondrial patterning during development"

**Supplementary Information:**

Figures S1 to S6
Table S1
Legends for Movies S1 to S6
Movies S1 to S6

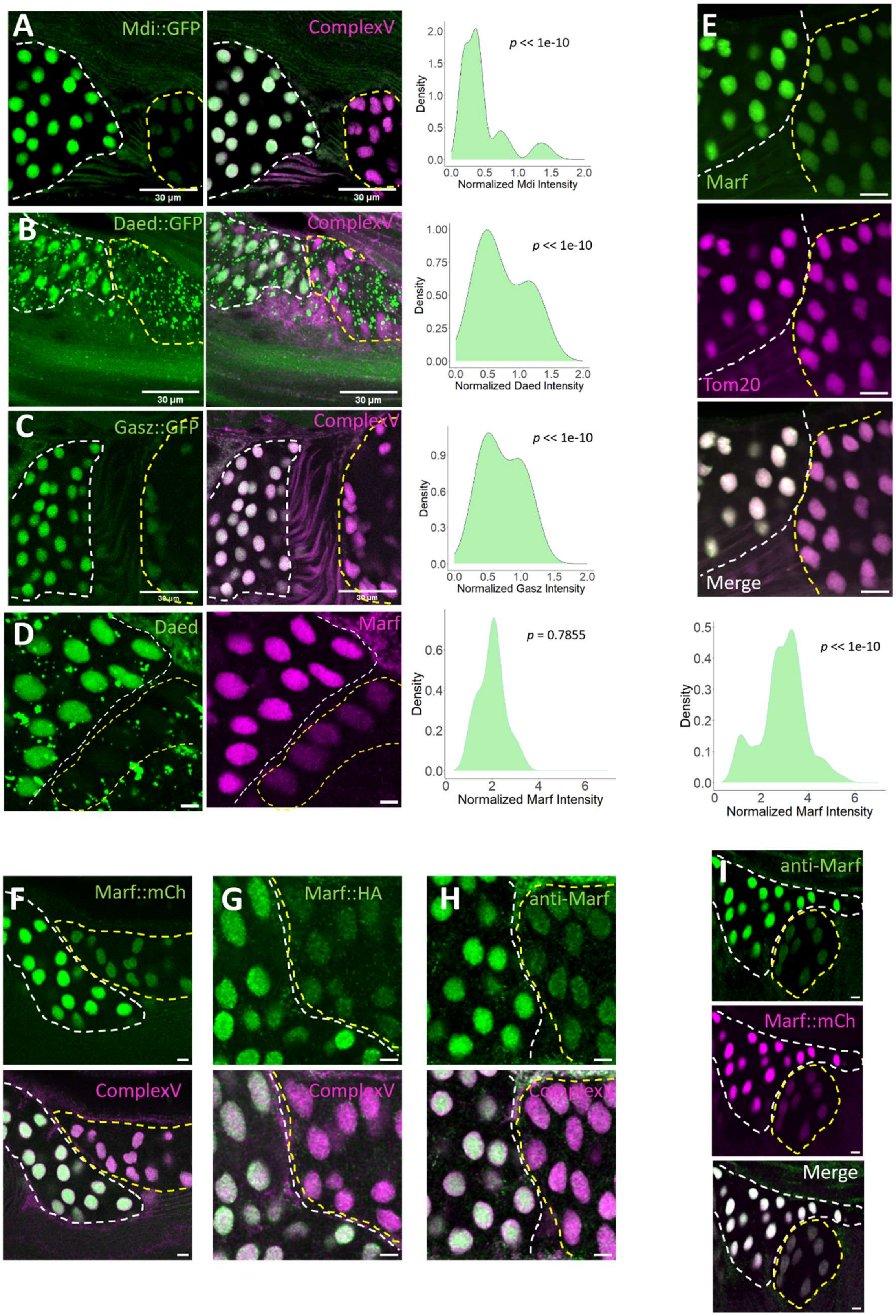
A
Mdi::GFP
ComplexV
30 µm
30 µm
p << 1e-10
Density
Normalized Mdi Intensity
B
Daed::GFP
ComplexV
30 µm
30 µm
p << 1e-10
Normalized Daed Intensity
C
Gasz::GFP
ComplexV
30 µm
30 µm
p << 1e-10
Normalized Gasz Intensity
D
Daed
Marf
p = 0.7855
Normalized Marf Intensity
E
Marf
Tom20
Merge
p << 1e-10
Normalized Marf Intensity
F
Marf::mCh
ComplexV
G
Marf::HA
ComplexV
H
anti-Marf
ComplexV
I
anti-Marf
Marf::mCh
Merge

**Fig. S1. (Associated with Fig. 2)**

(A) Two adjacent round spermatid cysts displaying high and low levels of Mdi in their nebenkern mitochondria. Density plot of Mdi intensity normalized to Complex V intensity (Right). n=307 nebenkerns from 3 testis. Nebenkerns show Mdi::GFP (green) and Complex V (magenta).

(B) Two adjacent round spermatid cysts displaying high and low levels of Daed in their nebenkern mitochondria. Density plot of Daed intensity normalized to Complex V intensity (Right). n=230 nebenkerns from 7 testis. Nebenkerns show Daed::GFP (green) and Complex V (magenta).

(C) Two adjacent round spermatid cysts displaying high and low levels of Gasz in their nebenkern mitochondria. Density plot of Gasz intensity normalized to Complex V intensity (Right). n=168 nebenkerns from 4 testis. Nebenkerns show Gasz::GFP (green) and Complex V (magenta).

(D) Two adjacent round spermatid cysts displaying high and low levels of Marf and Daed in their nebenkern mitochondria. Density plot of Marf intensity normalized to Daed intensity (Right). n=301 nebenkerns from 5 testis. Nebenkerns show Daed::GFP (green) and Marf stained with Marf antibody (magenta).

(E) Two adjacent round spermatid cysts displaying high and low levels of Marf with uniform Tom20 levels in their nebenkern mitochondria. Density plot of Marf intensity normalized to Tom20 intensity (Bottom). n=284 nebenkerns from 5 testis. Nebenkerns shows Marf::GFP (green) and Tom20::mCherry (magenta).

p-values<0.05 indicate a significant deviation from unimodality, estimated by Hartigans' dip test (A-E).

(F-H) Two adjacent cysts displaying high and low levels of Marf (green) visualized with Marf::mCherry (F), Marf::HA (G), and anti-Marf antibody (H). Complex V (magenta) is used to stain mitochondria.

(I) Marf stained with anti-Marf antibody (green) and Marf::mCherry (red). Fluorescent signals from antibody and mCherry fluorescence show good correlation, as evident by their faithful replication of high and low levels of Marf in the two adjacent cysts. The scale bars indicate 5μm.

.

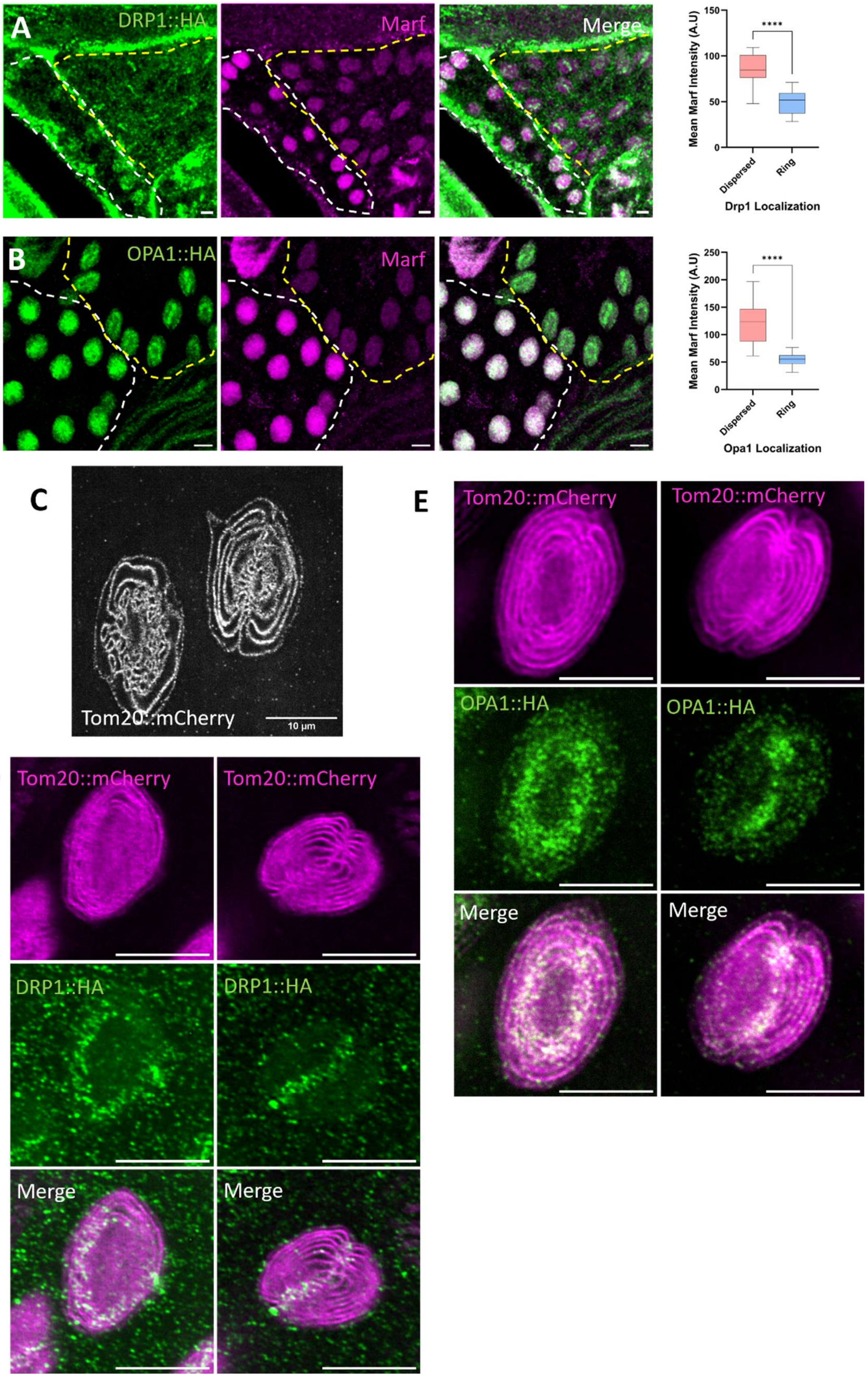
A
DRP1::HA
Marf
Merge
Mean Marf Intensity (A.U)
150
100
50
0
****
Dispersed
Ring
Drp1 Localization
B
OPA1::HA
Marf
Merge
Mean Marf Intensity (A.U)
250
200
150
100
50
0
****
Dispersed
Ring
Opa1 Localization
C
Tom20::mCherry
10 µm
D
Tom20::mCherry
Tom20::mCherry
DRP1::HA
DRP1::HA
Merge
Merge
E
Tom20::mCherry
Tom20::mCherry
OPA1::HA
OPA1::HA
Merge
Merge

**Fig. S2. (Associated with Fig. 2)**

(A) DRP1 localization in high- and low-Marf-containing nebenkerns. Quantification of Mean Marf intensity in DRP1 populations showing distinct localization patterns (Right). n=256 nebenkerns from 4 testis. Marf::GFP in magenta, DRP1::HA in green.

(B) OPA1 localization in high- and low-Marf-containing nebenkerns. Quantification of Mean marf intensity in OPA1 populations showing distinct localization patterns (Right). n=237 nebenkerns from 6 testis. Marf::GFP in magenta, OPA1::HA in green.

The error bar represents the standard error of the mean. An unpaired two-tailed t-test with Welch's correction was used to assess significance (A-B).

(C) Expansion microscopic image of two nebenkerns. The orientations of the nebenkerns reveal localized constrictions along the midplane of the nebenkern that looks like a ring (left) or like a straight line (right). Mitochondria marked with Tom20::mCherry

(D-E) SoRA super-resolution microscopy images of DRP1 punctae (D) and OPA1 punctae (E) in late nebenkerns. Depending upon the orientation of the nebenkern, DRP1 (D) and OPA1 (E) look like a ring (left) or a straight line (right), bifurcating the nebenkern. Mitochondria marked with Tom20::mCherry (magenta). DRP1::HA (D) and OPA1::HA (E) in green.

The scale bars indicate 5 µm unless otherwise noted.

.

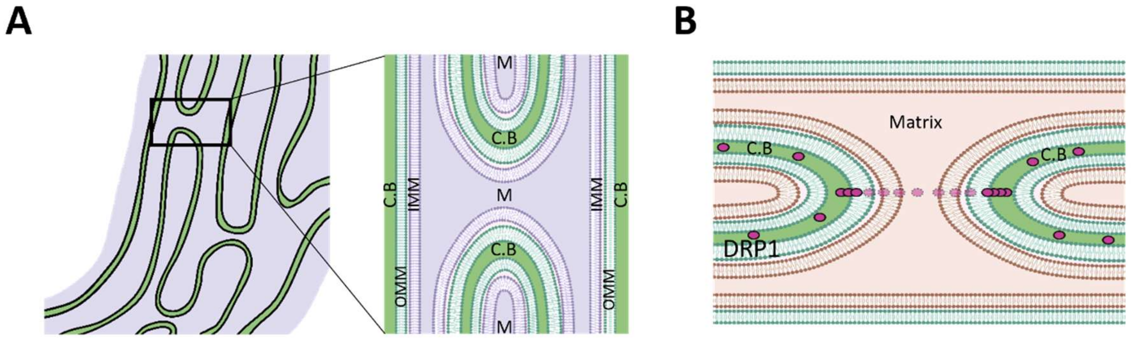


**Fig. S3. (Associated with Fig 3)**

(A) 2-D Schematic of inter-whorl connections tubulating the sheets. Inset shows the cytoplasmic bridges between two sets of OMM and IMM that make up each whorl. C.B - cytoplasmic bridges. M- matrix.

(B) Proposed model for geometry-determined DRP1 localization, resulting in DRP1-ring formation in the nebenkern midplane. Inter-whorl connections, such as in [A(inset)] tubulate nebenkern sheets. DRP1, which can pass through the cytoplasmic bridge, recognizes these tubulated regions and is stabilized in these regions. The localized tubulations along the nebenkern midplane thus restrict DRP1 stabilization to help bisect the nebenkern.

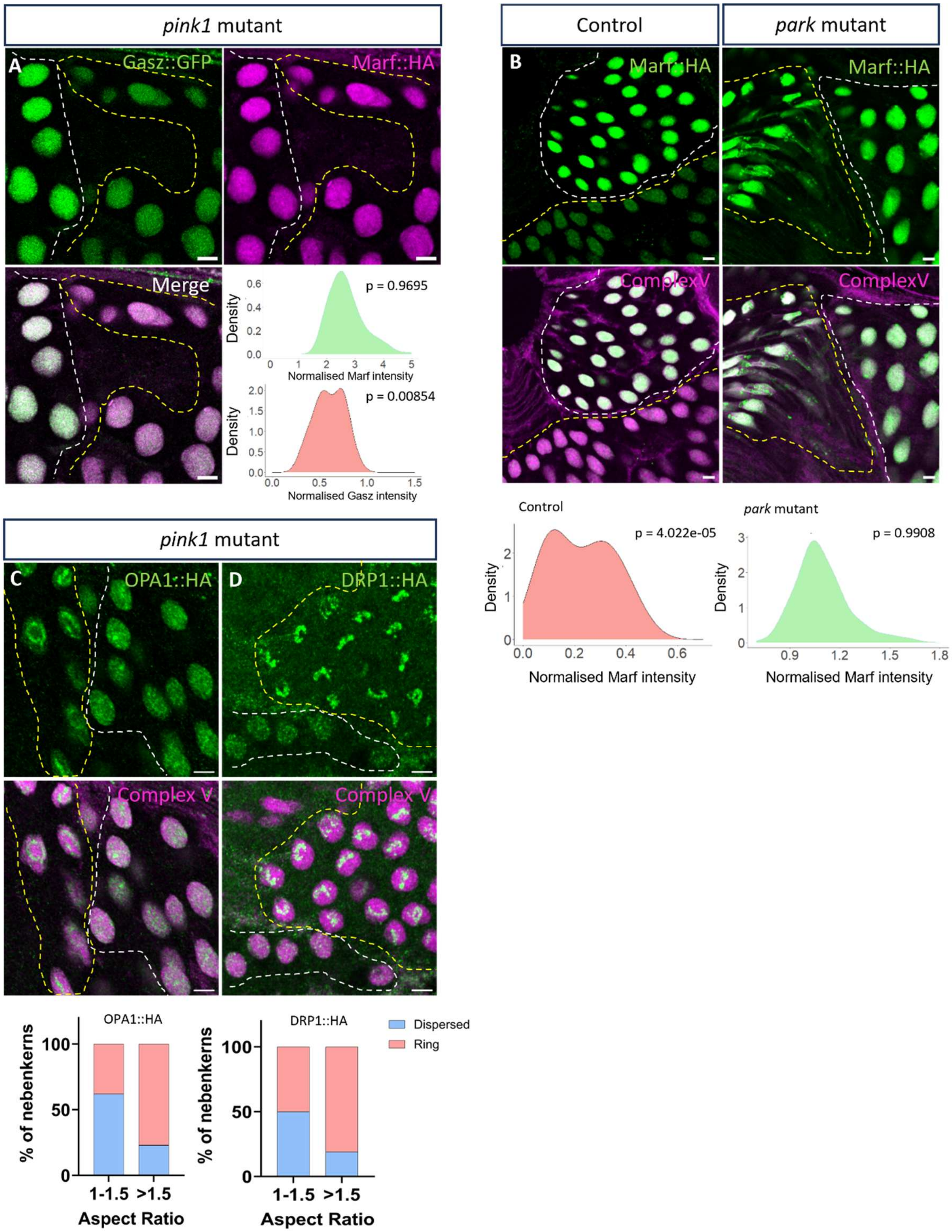
pink1 mutant
A
Gasz::GFP
Marf::HA
Merge
p = 0.9695
Density
Normalised Marf intensity
p = 0.00854
Density
Normalised Gasz intensity
Control
park mutant
B
Marf::HA
Marf::HA
ComplexV
ComplexV
Control
p = 4.022e-05
Density
Normalised Marf intensity
park mutant
p = 0.9908
Density
Normalised Marf intensity
pink1 mutant
C
OPA1::HA
D
DRP1::HA
Complex V
Complex V
OPA1::HA
% of nebenkerns
1-1.5
>1.5
Aspect Ratio
DRP1::HA
% of nebenkerns
1-1.5
>1.5
Aspect Ratio
Dispersed
Ring

**Fig. S4. (Associated with Fig 4)**

(A) Marf and Gasz levels in *pink1* mutants. Gasz levels (Top-Left) are downregulated in the nebenkerns of two adjacent cysts, while Marf levels (Top-Right) are unchanged. Nebenkerns show Gasz::GFP (green) and Marf (magenta). Density profiles (Bottom-Right) of Gasz, normalized to Complex V, show a multimodal distribution, while Marf, normalized to Complex V, shows a unimodal distribution. n=322 nebenkerns from 4 testis.

(B) Marf levels in control (*park[21],Marf::HA/+)* and *park (park21,Marf::HA/park[13])* mutants. Density profile of Marf normalized to ComplexV in control (Left) and *park* mutants (Right). n=263 nebenkerns from 5 testis in control and n=367 nebenkerns from 6 testis in mutant. Marf in green. ComplexV in magenta.

p-value<0.05 indicates a significant deviation from unimodality. Unimodality is estimated by Hartigans' dip test (A-B).

(C) OPA1 localization in early and late nebenkerns in *pink1* mutants. Quantification of OPA1 localization during nebenkern elongation. n=294 nebenkerns from 11 testis. OPA1::HA in green. ComplexV in magenta.

(D) DRP1 localization in early and late nebenkerns in *pink1* mutants. Quantification of DRP1 localization during nebenkern elongation. n=233 nebenkerns from 8 testis. DRP1::HA in green. ComplexV in magenta.

Chi-Squared analysis with Yates' correction shows a significant association of OPA1/DRP1 localization change with an increase in aspect ratio of nebenkerns (p-value: <0.0001) (C-D).

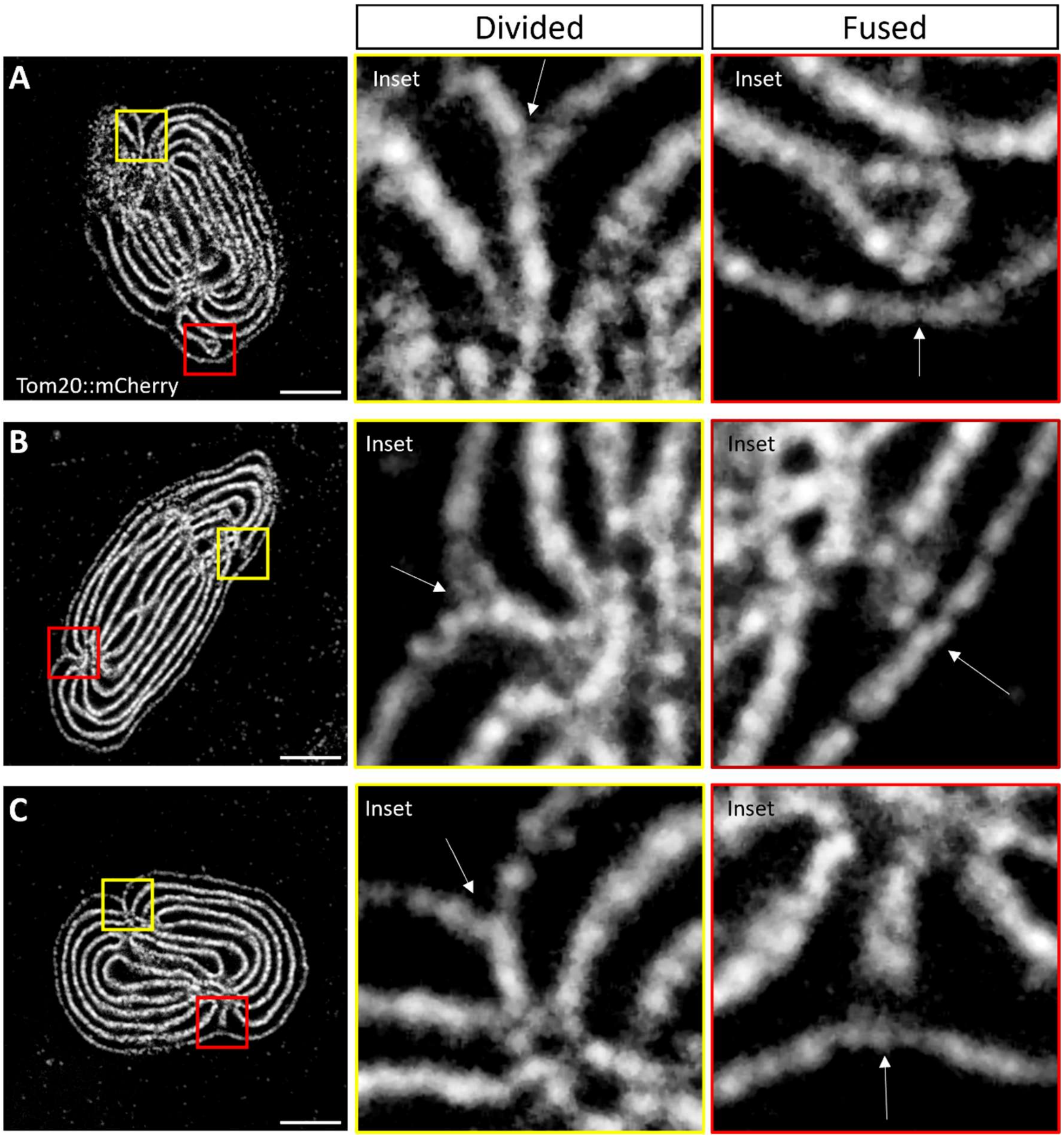


**Fig.S5. (Associated with Fig 4)**

(A-C) Examples of early-elongating *pink1* mutant nebenkerns, marked by Tom20::mCherry, showing fused outer whorls at one end and a divided state at the other. Yellow insets show a divided state. Red insets show the fused state.

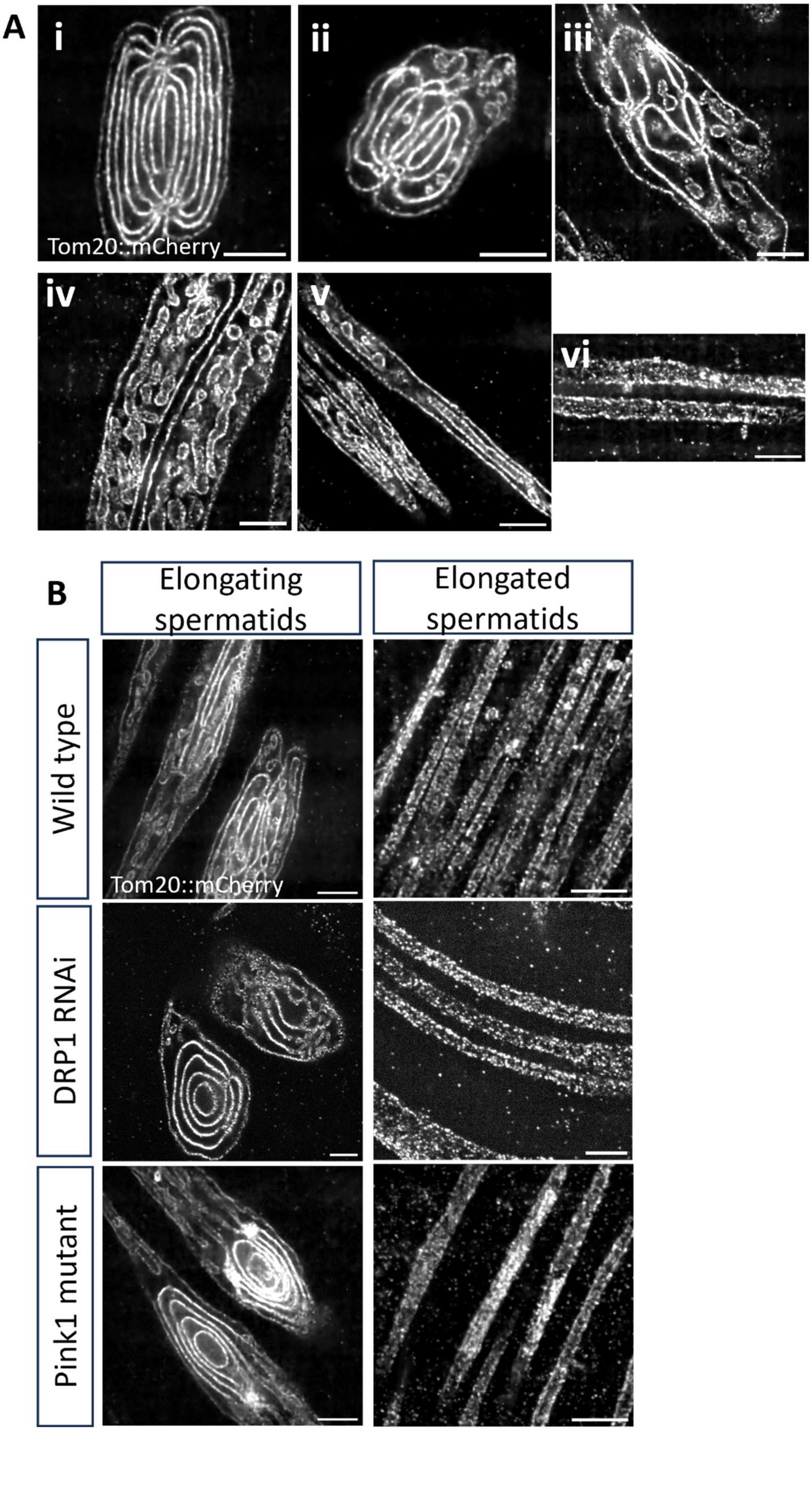

A
i
ii
iii
Tom20::mCherry
iv
v
vi
B
Elongating spermatids
Elongated spermatids
Wild type
Tom20::mCherry
DRP1 RNAi
Pink1 mutant

**Fig. S6. (Associated with Figs. 5 and 6)**

(A) Expansion microscopy images of mitochondria in different stages of elongation, from late nebenkerns (i) through fully elongated mitochondria derivatives (vi). MDiVs are absent in (i) and (vi); however, they increase steadily in the intermediary stages, while the number of whorls decreases (ii-v).

(B) Expansion microscopy images of elongating (Left) and elongated (Right) spermatids in wild-type (Top), DRP1 knockdown (Middle), and *pink1* mutants (Bottom). Elongating mitochondrial derivatives in all three conditions shows multiple whorls of membranes within the outer whorl, which are resolved during elongation, forming a tubular structure.

Scale bars are 5µm. Mitochondria are marked by Tom20::mCherry.

**Table S1.** A targeted protein expression screen of mitochondrial proteins in *Drosophila* spermatogenesis.

| Protein name | Mitotic stage | Meiotic stage | Nebenkern stage | Elongating stage |
|---|---|---|---|---|
| **Fission/Fusion** | | | | |
| **Marf/Mitofusin (Tagged line and antibody)** | **Uniformly present** | **Uniformly present** | **Two distinct levels: High and Low** | **Downregulated** |
| **OPA1 (tagged line)** | **Uniformly present** | **Uniformly Present** | **Two distinct localizations: Uniform and Ring-like** | **Uniformly present** |
| **DRP1 (Tagged line and antibody)** | **Predominantly cytoplasmic** | **Predominantly cytoplasmic** | **High mitochondrial localization with two distinct localizations: Uniform and Ring-like** | **Predominantly cytoplasmic** |
| **Quality control** | | | | |
| **PINK1 (Tagged line)** | **Low** | **Low** | **Highly upregulated** | **Comparable to nebenkern stages** |
| **HSP60 (antibody)** | **Low** | **Low** | **Highly upregulated** | **Comparable to nebenkern stages** |
| **piRNA biogenesis** | | | | |
| **Daed (Tagged line)** | **Uniformly present** | **Uniformly present** | **Two distinct levels: High and Low** | **Downregulated** |
| **Gasz (Tagged line)** | **Uniformly present** | **Uniformly present** | **Two distinct levels: High and Low** | **Downregulated** |

**Table S1. (Cont.)**

| Protein name | Mitotic stage | Meiotic stage | Nebenkern stage | Elongating stage |
|---|---|---|---|---|
| Biogenesis/Activity | | | | |
| Mdi (Tagged line) | Uniformly present | Uniformly present | Two distinct levels: High and Low | Downregulated |
| ATP5alpha (Antibody) | Uniformly present | Subtle downregulation | Levels comparable to downregulated meiotic stages | Levels comparable to downregulated meiotic stages |
| mtSSB (Tagged line) | Uniformly present | Low | Low | Low |
| Tfam (Tagged line) | High - Punctate | Low-Punctate | Low-Punctate | High-Punctate |
| Tamas (Tagged line) | Uniformly present | Low | Uniformly present | Uniformly present |
| Protein import | | | | |
| Tom20 (Tagged line) | Uniformly present | Uniformly present | Uniformly present | Uniformly present |

**Movie S1 (separate file).** 3D projection of OPA1 rings encircling maturing nebenkerns. Mitochondria marked ComplexV (red), and OPA1 marked with OPA1::HA (green). Scale bar 5 µm.

**Movie S2 (separate file).** 3D projection of DRP1 rings encircling maturing nebenkerns. Mitochondria marked ComplexV (red) and DRP1 marked with DRP1::HA (green). Scale bar 5 µm.

**Movie S3 (separate file).** Volumetric rendering of a late nebenkern after expansion microscopy. Clipping along the z-direction reveals the bifurcation and membrane tubulation along the middle groove. The mitochondrial membrane is marked with Tom20::mCherry. Scale bar 5 µm.

**Movie S4 (separate file).** Volumetric rendering of an isolated inter-whorl connection. Rotation about the x-axis reveals the tubular connection traversing a middle whorl, resulting in holes in both sheets. Clipping shows the constriction in the middle whorl. The mitochondrial membrane is marked with Tom20::mCherry. Scale bar 5 µm..

**Movie S5 (separate file).** Volumetric rendering of the elongating tip of a mitochondrial derivative after expansion. The tip shows several MDiVs filled with cytoplasmic contents. Mitochondrial membranes are marked with Tom20::mCherry (magenta) and cytoplasm with cytoplasmic GFP (green). Scale bar 5 µm.

**Movie S6 (separate file**). Volumetric rendering of an MDiV suspended within a mitochondrial derivative after expansion. Clipping shows a point of membrane continuity between the mitochondrial derivative and the MDiV, suggesting fusion. The mitochondrial membrane is marked with Tom20::mCherry. Scale bar 2 µ**m.**
